# BUB1B MITOTIC KINASE DRIVES THERAPY RESISTANT PROSTATE CANCER

**DOI:** 10.1101/2025.11.09.687448

**Authors:** MJ Martínez, C Blanco, N Peinetti, RLZ Lyles, MV Revuelta, KL Burnstein

## Abstract

Castration-resistant prostate cancer (CRPC) that progresses despite treatment with potent AR antagonists such as enzalutamide is a major clinical problem. Through an unbiased systems biology approach, we previously identified a therapeutically relevant seven gene set that drives CRPC. The mitotic checkpoint kinase, BUB1B (BUBR1) is a key member of this gene set, and we report here that BUB1B is a tractable and promising new therapeutic target in aggressive, treatment-resistant PC. We found that high BUB1B expression is correlated with PC progression and aggressiveness. In established CRPC cells, BUB1B depletion blocked cell proliferation through cell cycle arrest and mitosis delay. Ectopic expression of BUB1B, at levels found in CRPC, conferred castration-resistant growth of androgen-dependent PC cells *in vitro* and *in vivo*. We showed that BUB1B kinase activity was essential for CRPC progression, as only wild type (wt) BUB1B and neither of two kinase-dead mutants promoted castration-resistant growth of androgen-dependent PC cells. Rescue experiments with wt or kinase-dead mutants showed further that BUB1B kinase activity was also required to maintain proliferation of established CRPC cells. While persistent androgen receptor (AR) signaling is a mechanism of CRPC progression, BUB1B promotion of CRPC was not dependent on AR as assessed through AR knockdown. Consistent with an AR-bypass mechanism, ectopic expression of BUB1B rendered PC cells resistant to enzalutamide *in vitro* and *in vivo*. Our data points to BUB1B as a key driver of CRPC progression and enzalutamide resistance and suggests that targeting BUB1B kinase is a promising therapeutic approach for lethal, treatment-resistant disease.

**Statement of significance:** We document a novel role for BUB1B kinase as a critical driver of castration- and enzalutamide-resistant prostate cancer, highlighting the therapeutic potential of BUB1B kinase inhibition to overcome lethal treatment-resistant disease.

## Introduction

Androgen deprivation therapy (ADT) is the standard of care for patients with advanced or high-risk prostate cancer (PC); however, after 2-3 years most patients relapse and develop incurable castration resistant prostate cancer (CRPC)^1–4^. Second generation androgen receptor (AR) antagonists or AR signaling inhibitors (ARSIs), such as enzalutamide and abiraterone, improve survival of men with CRPC by a few months^5–7^. A variety of molecular mechanisms can promote CPRC progression, including AR-dependent and -independent pathways. AR-dependent mechanisms include increased *AR* levels, expression of constitutively active AR variants (AR-Vs), and local synthesis of androgens^8^. AR-independent pathways include increased expression and activity of glucocorticoid receptors (GR), activation of signaling pathways (e.g. AKT/mTOR) as well as cellular plasticity leading to neuroendocrine transdifferentiation^9,10^.

We previously reported an unbiased, integrated systems biology approach to define CRPC vulnerabilities that arise through ligand-independent AR signaling. This effort identified a seven gene set driving CRPC with several actionable therapeutic targets^11^. This gene set is broadly overexpressed across CRPC and strongly predicts poor disease-free and overall survival in PC patient cohorts. The mitotic serine/ threonine kinase, BUB1B (BUBR1), is a central member of the seven gene set. BUB1B is a core component of the spindle assembly checkpoint (SAC), which is a surveillance pathway that blocks anaphase progression until all sister chromatids are correctly positioned thereby avoiding mitotic errors(^Reviewed in 12–14^). Although previously characterized as a pseudokinase^15^, both BUB1B autophosphorylation and BUB1B-mediated phosphorylation of CENP-E ensure proper microtubule-kinetochore attachment in human cells^16–19^. In addition to our report^11^, bioinformatic analyses of patient datasets have identified high BUB1B mRNA levels to be associated with aggressiveness and progression in PC patients^20–22^. High BUB1B protein levels in patient tissue samples have also been proposed to be prognostic and associated with higher Gleason scores in PC^23,24^. While BUB1B knockdown decreases PC cell proliferation^11,24,25^, it remains unknown whether BUB1B and/or its catalytic function is required for progression to, and maintenance of CRPC and as well as enzalutamide resistance.

Here we show that BUB1B mRNA and protein are up-regulated in CRPC cell lines, advanced PC patient samples and CRPC patient-derived xenografts (PDXs). Elevated BUB1B is associated with worse prognosis. Moreover, BUB1B knockdown halted CRPC proliferation by cell cycle inhibition and mitosis delay. BUB1B ectopic expression, at levels comparable to CRPC, was sufficient to convert androgen dependent (AD) PC cells to enzalutamide-resistant CRPC *in vitro* and *in vivo*. We demonstrated that BUB1B kinase activity was required for both progression to and maintenance of CRPC thereby positioning BUB1B kinase as a promising therapeutic target for CRPC.

## Materials and Methods

### Analysis of human samples datasets

The TCGA-PRAD was downloaded from Xena Functional Genomics Explorer (http://xena.ucsc.edu/), MSKCC (Cancer cell, 2010) and SU2C (Cell, 2019) data sets were accessed and downloaded from cBioportal (cbioportal.org). The GSE9476726 and GSE3226927 data sets were downloaded from GEO portal (https://www.ncbi.nlm.nih.gov/gds). The BUB1B expression values for all datasets were presented as log2-transformed signal intensity ratios. To build Kaplan-Meier curves, patient samples were stratified in quartiles based on BUB1B expression and disease-free curves were built comparing patients in the top quartile (high BUB1B) with patients in the bottom quartile (low BUB1B).

### Cell culture and reagents

Human PC cell lines RWPE-1, LNCaP, 22Rv1 (CRL-2505), DU145 and PC3 were obtained from [American Type Culture Collection (ATCC)]. VCaP were a gift from Dr Pienta, CWR-R1 were a gift from Dr Wilson, C4-2B were a gifts from Dr Connor Lynch. GP2-293 were purchased from Clontech. All cell lines were negative for mycoplasma, bacteria, and fungi contamination. LNCaP, 22Rv1, and PC3 cells were cultured in RPMI (Corning, 15-040-CV) supplemented with Penicillin/ Streptomycin Solution 100X (100ml) (Corning, #30-002-CI), 200 mM l-glutamine (Corning, # 25-005-CL), and 10% fetal bovine serum (FBS) (Hyclone). C4-2B cells were cultured in DMEM (Corning, #10-013-CV), supplemented as detailed above. VCaP cells were cultured in DMEM high glucose Glutamax (Thermo, 10569-044), antibiotic antimycotic solution (Gibco 15240-062) and 10% FBS. For androgen depletion, cells were washed with warm PBS and incubated twice with unsupplemented media for an hour, and media was replaced with 2% (LNCaP and CRPC cells) or 10% (VCaP) CSS. Cell cultures were maintained at 37°C in a humidified atmosphere of 5% CO_2_. Cells were selected with Puromycin Dihydrochloride (Thermo Fisher, A1113803) or G418 solution (Millipore-Sigma, G418-RO). For drug treatment experiments, cells were treated with enzalutamide (MDV3100, Cayman Chemical), GSK-923295 (HY-10299, MedChem) or mifepristone (Tocris, 1479).

### Plasmids

The pLKO.1-shBUB1B plasmids against the 3’UTR (TRCN0000197142 and TRCN0000000461) or the coding region (CDC) (TRCN0000194741) were purchased from Sigma-Aldrich. The pQCXIN retroviral vector was purchased from BD Biosciences. pLKO.1-shGFP plasmid was provided by Dr. Priyamvada Rai (University of Miami). The viral packaging plasmids, VSV-G and Δ8.2, were purchased from Clontech. For overexpression experiments, pCMV-6 BUB1B wild type was purchased from Origene (RC205844), and pLAP-BUB1B D882B (114062) and K795R (114060) were purchased from Addgene. Then, BUB1B wild type, D882N and K795R were subclone into pQCXIN plasmid. Cells transduced with pLKO.1 plasmid were selected in 2.5 µg/ml of puromycin for 2 to 3 days. pQCXIN transduced cells were selected with 0.75-1 mg/mL G418 (Millipore-Sigma, #04727878001, G418-RO) for 7 to 10 days.

### *In vitro* cell proliferation assay

For BUB1B knockdown experiments, 20,000 cells per well (LNCaP, 22Rv1 and PC3), or 40,000 cells per well (C4-2B) were seeded in 24 well plates. The following day, the media of LNCaP ADPC cells was replaced with 2% FBS media (time = 0); CRPC cells were androgen depleted, and media was replaced with 2% CSS media (time = 0). For overexpression experiments, 22Rv1 and LNCaP EV and BUB1B wild type, D882N or K795R cells were androgen depleted (as described above), and 20,000 cells were seeded in 2% CSS media in 24 well plates. For VCaP EV and BUB1B wild type, D882N or K795R, 100,000 cells were seeded in 24 well plates. The following day, cells were androgen depleted as previously described, and media was replaced with 10% CSS media. For rescue experiments, 22Rv1 and LNCaP EV and BUB1B wild type, D882N or K795R cells were transfected after androgen depletion using siBUB1B (against 3’ UTR) or siControl (Dharmacon). For drug treatment experiments, cells were androgen depleted and media was replaced with 2% (LNCaP) or 10% (VCaP) CSS. For enzalutamide treatment, cells were treated with 10 µM of drug or vehicle on day 0. For GSK-923295 experiments, cells were treated with 30, 60, and 120 nM for 7 days. To assess combination therapy, cells were treated with enzalutamide (10, 20, and 30 µM), GSK-923295 (10, 20, and 30 nM) or their combination. At different time points (0-8 days), cells were trypsinized, centrifuged for 10 min at 5,500 rpm and resuspended in 20 µL of media. 10 µL of the cell suspension was mixed with 10 µL of trypan blue (Gibco), and live cells were counted using the Countess II (Thermo Fisher). Data represents at least three independent experiments performed in quadruplicate.

### Soft agar assay

Soft agar assays were performed as previously described^28,29^. Briefly, LNCaP EV and BUB1B cells were androgen depleted and 200,000 cells were seeded in 2% CSS soft agar plates. Fresh media was added once a week. Plates were incubated for 3 weeks. Colonies were stained with 0.005% crystal violet, imaged under microscope and colonies were counted using Image J.

### Live cell imaging for caspase -3/-7 activity assay and mitosis progression

For caspase -3/-7 assay, cells were androgen depleted (see above) 24 h after seeding and transfected with 1% (v/v) IncuCyte Caspase-3/-7 Apoptosis Assay Reagent (Essen Bioscience). Incucyte Zoom software was used to analyze the phase and green-fluorescent images. Cells were cultured in 10% CSS and imaged every 4 h in the Incucyte Zoom System (Essen Bioscience) at 10X magnification. For mitosis progression, 20,000 cells per well were seeded in 24 well plates. The following day, cells were androgen depleted and imaged every 20 minutes for 48 h using the Incucyte Zoom System (Essen BioScience) at 20X magnification. Individual cells (50 cells per group) were tracked from the beginning to end of mitosis, and mitosis duration was calculated.

### Cell cycle analysis

Cells were seeded in 10 cm^2^ dishes and incubated overnight at 37°C in a humidified atmosphere of 5% CO_2_. The following day, shBUB1B or shGFP cells were trypsinized, pelleted down, and washed twice with cold PBS. Pellet was re-suspended in 200 µL of PBS + 2 mM EDTA and fixed by adding 600 µL of cold 70% ethanol followed by overnight incubation at 4°C. Before flow cytometry analysis, cells were centrifuged at 4°C at 1000 rpm for 5 min and re-suspended in 500 µL of PBS + 2 mM EDTA. 10 µL heat inactivated RNAse A (10 mg/mL) and 20 µL of propidium iodide (Sigma-Aldrich) were added. Cells were incubated in a water-bath at 37°C for 30 min and analyzed using a flow-cytometer.

### Immunoblotting

200,000- 600,000 cells / well were seeded in 6 well plates and incubated overnight. The following day cells were androgen depleted, and media was replaced with fresh 2% CSS media. Cells were harvested in RIPA buffer (Rockland) with protease inhibitor cocktail (Thermo Scientific). Protein samples were denatured in Laemmli buffer with β-mercaptoethanol and boiled at 95°C for 10 min. Western blots were performed as previously described^29,30^. Anti-Cleaved PARP (9541), anti-GR (3660T), anti-AR-V7 (68492), anti-rabbit (7074s), and anti-mouse (7076s) antibodies were purchased from Cell Signaling Technology. Anti-Actin (sc-1616), anti-Cyclin A (sc-751), anti-CDK-1 (sc-54) and anti-goat (sc-2354) were purchased from Santa Cruz Biotechnology. Anti-synaptophysin antibody [YE269] (ab32127) and anti-BUB1B (ab54894) antibody was purchased from Abcam. Anti-AR (06-680) antibody was purchased from Millipore. Anti-HSP90 (610419) was purchased from BD Biosciences. GAPDH was purchased from Invitrogen (MA5-15738).

### Subcutaneous xenografts

Experiments with mice were conducted under an approved IACUC (Institutional Animal Care and Use Committee) protocol with the University of Miami and adhered to the NIH (National Institutes of Health) Guide for the Care and Use of Laboratory Animals. Pre-castrated SCID male mice (Envigo, C.B-17/IcrHsd-Prkdc-scid), 5 to 6 weeks old were used. 2×10^6^ LNCaP EV (n = 10) or BUB1B (n=29) cells were injected into the right hind flank resuspended in DPBS and Matrigel (Corning, 356234). Mice were weighed once a week. Tumor volume was measured by caliper twice a week, and volume was calculated using the equation: Length × Width × Depth × 0.52. The endpoint was defined when tumors reached 500 mm^3^. Animals were euthanized by CO_2_ and cervical dislocation; blood and tumors were collected, weighed and imaged. Part of the tissue collected from EV or BUB1B tumors was used to generate cell lines. Briefly, 1 ml of P XIV solution [25 mg P XIV (Sigma-Aldrich, P5147), 50 mL DMEM, antibiotic-antimycotic (100x) (Thermo Fisher, 15240062), 50 µl Gentamicin (Sigma-Aldrich, G1397) and 50 µl DNAseI (Sigma-Aldrich, DN25)] was added to the tumor tissue, tissue was homogenized and was incubated at 37°C for 30. Pellet was resuspended in 2 mL of 1X RBC lysis buffer (Biolegend, 420301) and incubated for 12 min on ice. Cells were washed twice with 20 mL cold PBS and resuspend in 10 mL RPMI 10% FBS. Two cell lines were generated, one derived from EV tumor (T_EV) and another derived from BUB1B tumor (T_BUB1B).

For enzalutamide treatment, pre-castrated SCID male mice (Envigo, C.B-17/IcrHsd-Prkdc-scid), 5 to 6 weeks old were used. 2×10^6^ LNCaP EV or T_BUB1B cells resuspended in DPBS and Matrigel (Corning, 356234) were injected into the right hind flanks (10 mice for EV and 29 mice for BUB1B). Mice were weighed once a week. Tumor volume was measured by caliper twice a week, and volume was calculated using the equation: Length × Width × Depth × 0.52. Once tumors reach 200 mm^3^, animals were randomly divided into two groups and treated daily with enzalutamide (n=9) (10 mg/kg; Cayman Chemical, 11596) or Vehicle (n=11) (0.3% Methylcellulose; SigmaAldrich, #M7140). Animals were dosed daily via oral gavage for 30-40 days. The endpoint was defined when tumors reached 1000 mm^3^. Animals were euthanized by CO_2_ and cervical dislocation; blood and tumors were collected, and tumors were weighed.

Circulating PSA (Prostate Specific Antigen) was measured by enzyme-linked immunosorbent assay (BioCheck Inc., #BC-1019) from blood plasma. Tumor tissue was homogenized, and protein was collected for western blot analysis of BUB1B levels in tumor samples.

### Immunohistochemistry

BUB1B protein was determined by immunohistochemistry (IHC) for tissue samples from AD and CR LuCaP PDXs pairs (LuCaP35, LuCaP77 and LuCaP96) and tumor micro arrays (TMAs) containing tissue from metastases [16 from bone and 17 from visceral tissue (liver, spleen, lung and lymph node)] collected from 10 patients or non-tumor samples (n=2). IHC was performed by the Cancer Modeling Shared Resource (CMSR, University of Miami) using Abcam BUB1B antibody (ab54894). Images were captured using the OlyVIA (Olympus Viewer for Imaging Applications) software and were quantified using Visiopharm software.

### Statistical analysis

Data were tested for normality (Shapiro–Wilk test) and homogeneity of variances (Levene’s test). When assumptions were met, data were tested for significance (P < 0.05) using a two-tailed Student’s t-test (two groups) or analysis of variance (ANOVA; three or more groups). Otherwise, non-parametric statistical analyses were used: Mann– Whitney’s test (two groups) and Kruskal–Wallis (unpaired, three or more groups) or Friedman (paired, three or more groups). Log-rank tests were used to test the significance of differences in Disease Free curves. Results are expressed as means ± SEM, and P < 0.05 was considered significant. Combinatory index (CI) was calculated using the CompuSyn software. Statistical analyses and graph plotting were carried out using GraphPad Prism software version 8.4 (www.graphpad.com). For the in vivo experiment, tumor volume over time between EV and BUB1B groups was evaluated using a mixed model with cubic effect. Differences were observed between groups at the end of the experiment (day 85). Analysis was performed by the Biostatistics & Bioinformatics Shared Resource (BBSR, University of Miami).

## Results

### 1. High levels of BUB1B correlate with aggressive PC

To determine whether BUB1B expression was higher in more advanced stages of the disease, we first stratified the SU2C and MSKCC data into BUB1B high- and low-expressing patients. Patients with metastatic CRPC (mCRPC) and high BUB1B expression (top quartile) had reduced disease-free survival and overall survival compared to low expression (bottom quartile) (Figure 1 a). Furthermore, BUB1B mRNA levels were higher in mCRPC patient tumors compared to localized (primary) PC and in PC compared to non-cancer in patient sample data sets (Figure 1 b). Similarly, BUB1B protein levels were elevated in three castration-resistant LuCaP patient derived xenograft (PDXs) (LuCaP35, LuCaP77 and LuCaP96) tissue compared to their AD counterparts^31^ (Figure 1 c). In human tissue microarrays (TMAs) containing samples from 10 patients with 16 bone and 17 visceral tissue (liver, spleen, lung and lymph node) metastases, we found that BUB1B protein levels were higher in metastatic CRPC tumor tissue compared to normal tissue (Figure 1 d). Like patient samples, BUB1B (mRNA and protein) was higher in established human PC and CRPC cell lines than in non-tumorigenic RWPE-1 prostate epithelial cells. Also, BUB1B mRNA and protein levels were higher in the castration-resistant derivative C42B compared to its AD parental line LNCaP (Figure 1 e and f).

**Figure 1.**
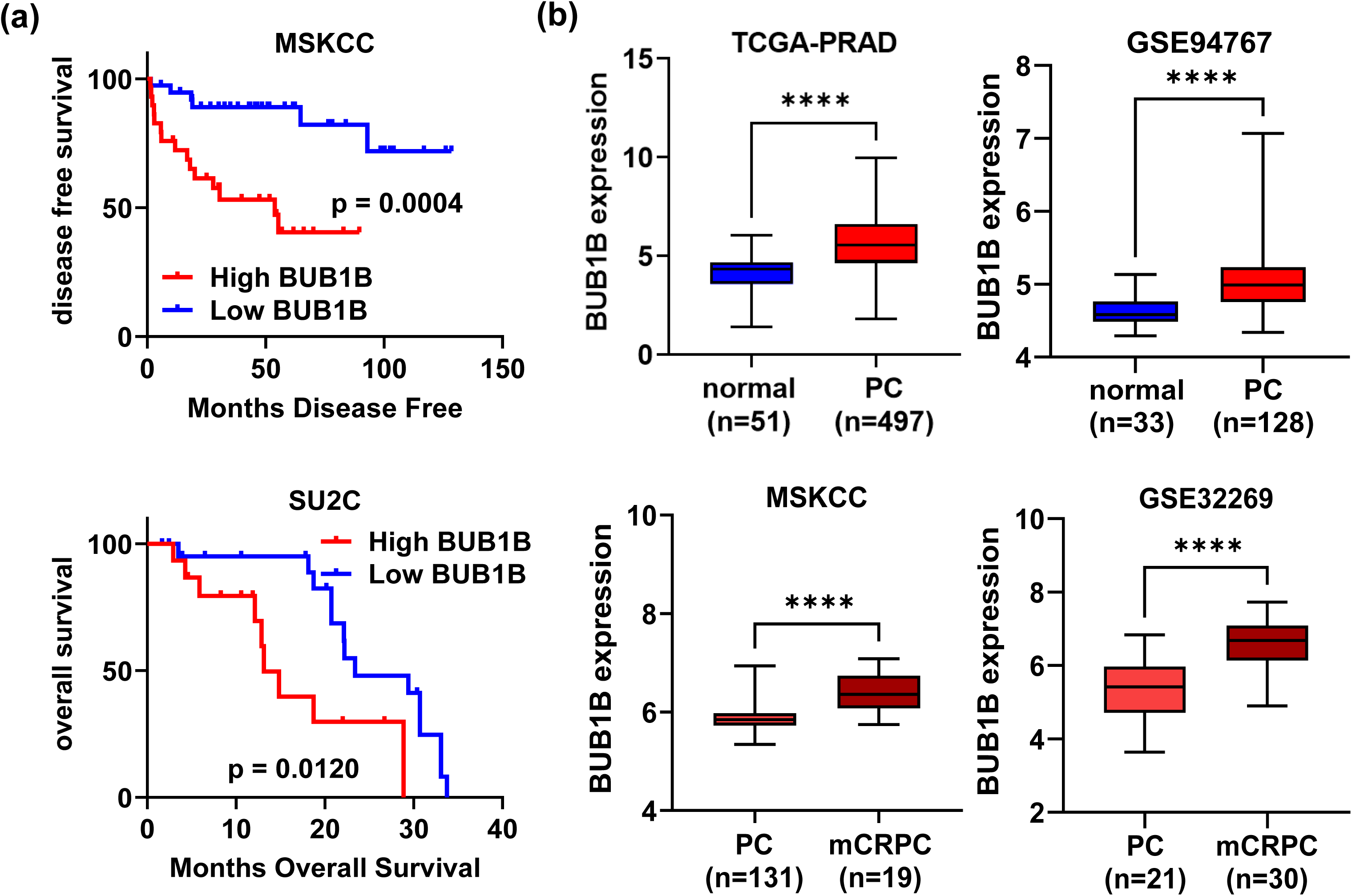
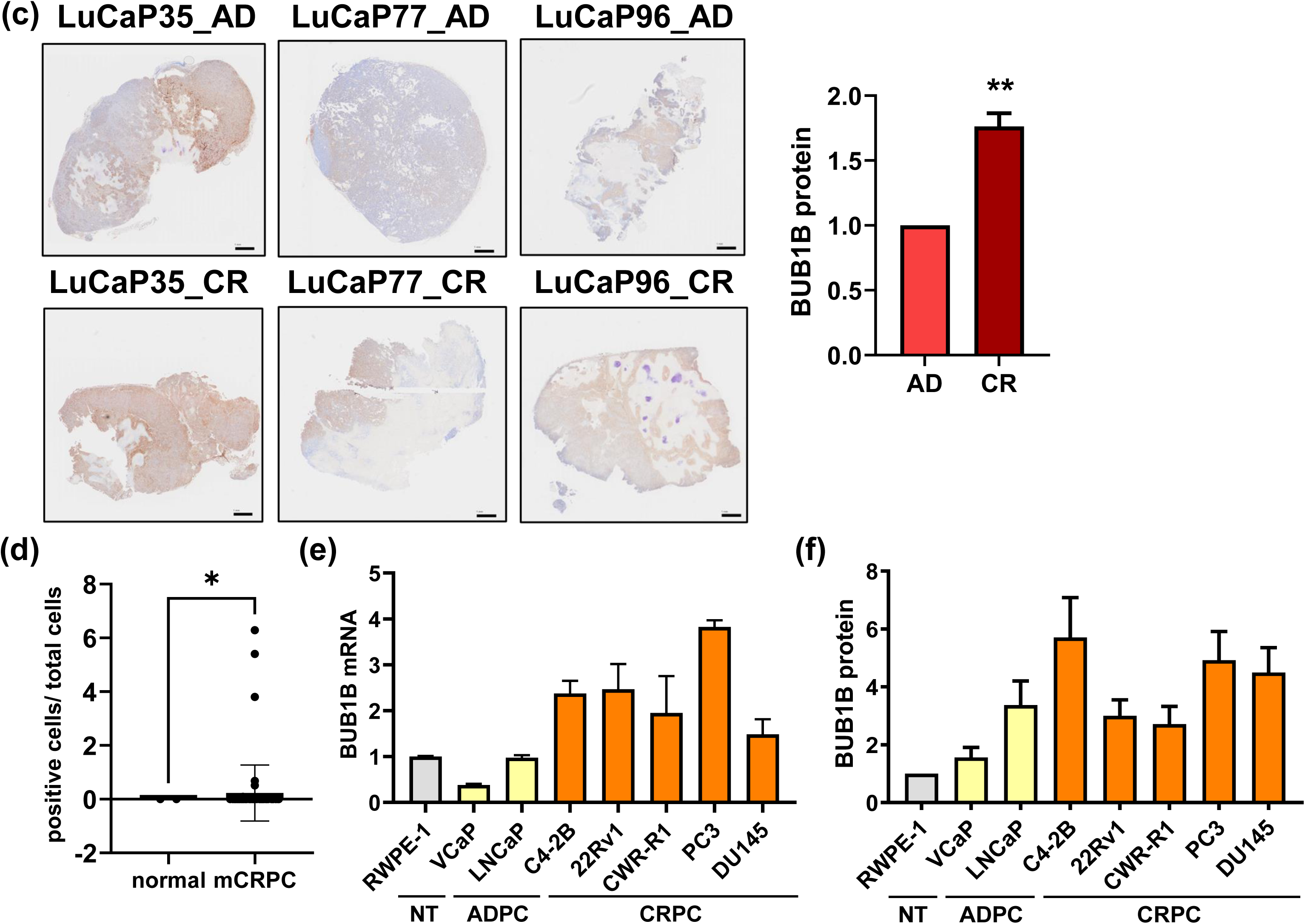
BUB1B expression is elevated in human PC and associated with poor prognosis. **(a)** Disease Free Survival or Overall Survival was evaluated in mCRPC patients with high BUB1B expression (top quartile) compared to patients with low BUB1B expression (bottom quartile) from (Cancer Cell, 2010) and SU2C RNA-seq data sets (downloaded from cBioportal) (log-rank test, ** p < 0.01, *** p < 0.001). **(b)** BUB1B mRNA expression was compared between normal or PC samples or between different disease stages in TCGA-PRAD (RNAseq data downloaded from http://xena.ucsc.edu/) MSKCC (Cancer Cell, 2010 from cBioportal), GSE94767, GSE32269, (microarray data from https://www.ncbi.nlm.nih.gov/gds) (student’s t test, **** p < 0.0001). BUB1B protein was evaluated by IHC in **(c)** three pairs of LuCaP androgen dependent (AD) and castration resistant (CR) patient derived xenografts (PDXs) (LuCaP35, LuCaP77 and LuCaP96) or **(d)** in TMAs from metastatic CRPC samples from 10 patients with 16 bone and 17 visceral tissue (liver, spleen, lung and lymph node) (n=33) and benign tissue (n=2). Graphs represent the quantification of BUB1B positive cells IHC for **(c)** LuCaP PDXs and **(d)** TMAs. (student’s t test, * p < 0.05, ** p < 0.01). BUB1B **(e)** mRNA and **(f)** protein levels were evaluated in RWPE-1 [non-tumorigenic (NT), grey], LNCaP, VCaP (ADPC, yellow), C4-2B, 22Rv1, CWR-R1, PC3 and DU145 (CRPC, orange) cells. Graphs represent the average ± SEM of two or three independent experiments performed in triplicate.

### 2. BUB1B depletion decreases proliferation and cell cycle progression of CRPC cells

To test the importance of BUB1B in CRPC proliferation, BUB1B was depleted using three different shRNA constructs [two targeting the 3’ UTR and one to the coding region (CDR)] in 22Rv1 CRPC cells. Although all three constructs decreased BUB1B expression and cell proliferation; the 3’ UTR#1 decreased growth of 22Rv1 cells to a greater extent and was therefore selected for further studies (Supplementary Figure 1 a). BUB1B knockdown completely halted proliferation of all CRPC cell lines tested including both AR expressing and AR negative lines while having more modest anti-proliferative effects on the ADPC cell line LNCaP (Figure 2 a). Growth inhibition following BUB1B depletion was accompanied by reduced CDK1 and Cyclin A levels and increased accumulation in G2/M phase specifically in CRPC cells (Figure 2 b and c, Supplementary Figure 1 b and c). Moreover, BUB1B depletion significantly prolonged mitosis duration (Figure 2 d). BUB1B knockdown did not induce caspase-dependent apoptosis, as evidenced by lack of increased cleaved PARP or Caspase -3/-7 activity (Supplementary Figure 1 d-f). These data show that BUB1B depletion decreased CRPC proliferation through inhibition of cell cycle and mitotic progression in line with its canonical role as a central regulator of the spindle assembly checkpoint.

**Figure 2.**
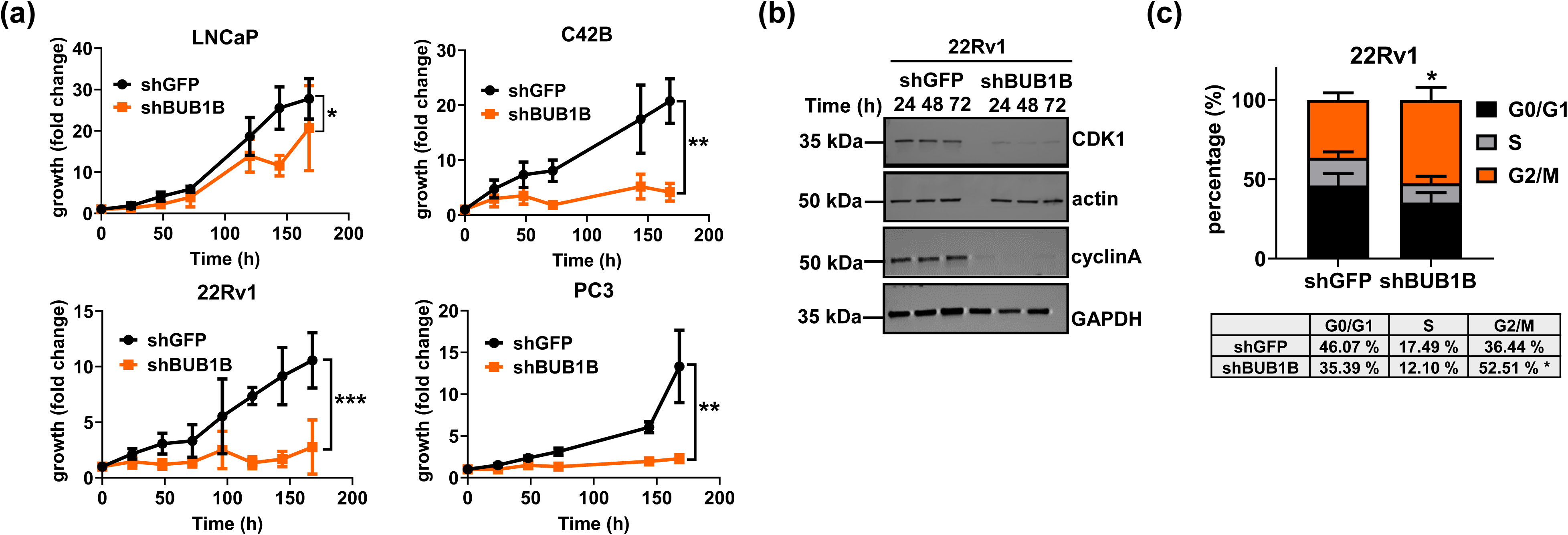
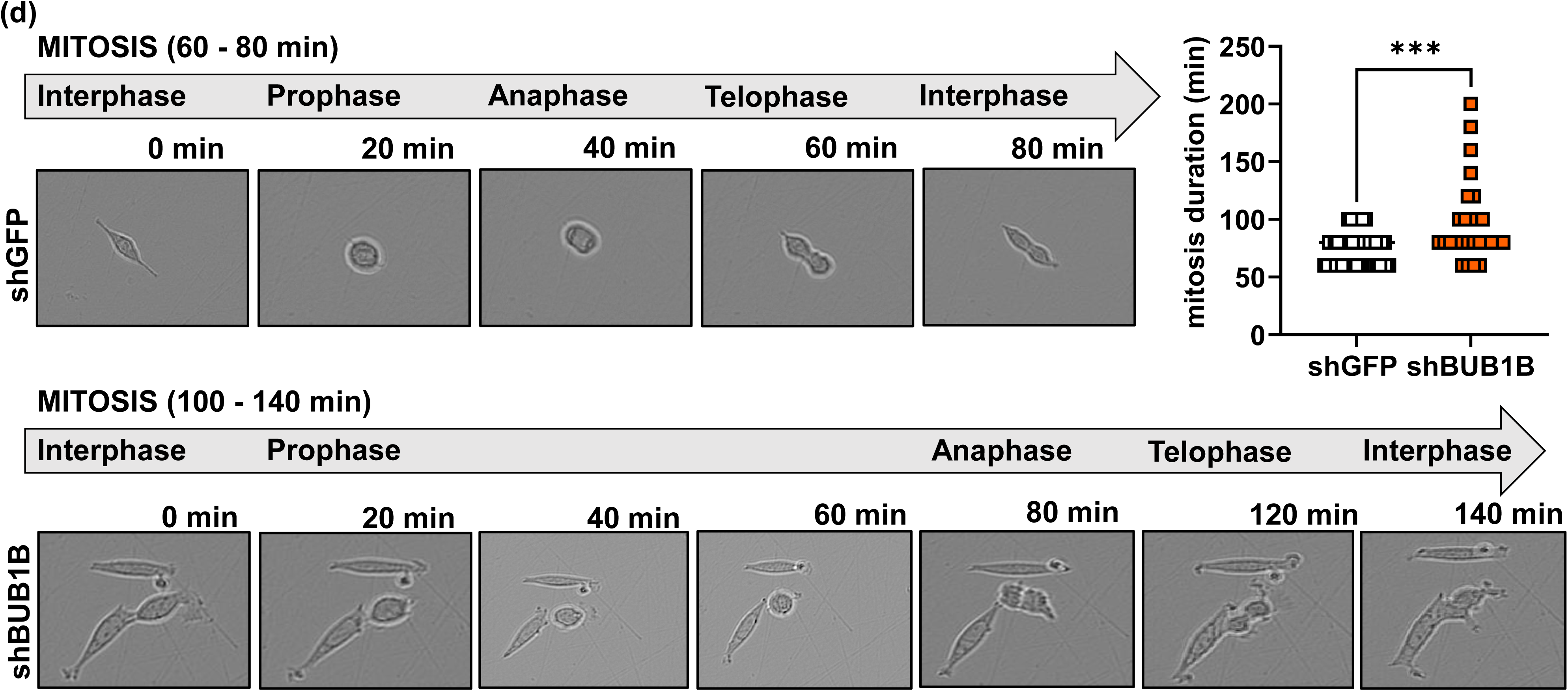
BUB1B depletion decreased CRPC growth through cell cycle inhibition. **(a)** Cells excluding trypan blue were counted following transduction with shBUB1B (3’UTR) or control (shGFP) in LNCaP (AR positive) ADPC cells, C4-2B (AR positive), 22Rv1 (AR/ AR-V7 positive) and PC3 (AR-negative) CRPC cells. Graphs represent the average of four or five independent experiments performed in quadruplicate ± SEM. The area under the curve was calculated for each cell line (Mann-Whitney test, * p < 0.05, *** p < 0.001). **(b)** CDK1 and cyclin A levels after BUB1B depletion in 22Rv1 by Western blot. Western blot quantification is shown in Supplementary Figure 1 b. **(c)** Cell cycle distribution was measured after BUB1B depletion (vs control shGFP) in 22Rv1 cells by flow cytometry of propidium iodide (PI) stained cells. Graphs represent distribution of the cell population across the cell cycle of the average ± SEM of four independent experiments (Mann-Whitney test, * p < 0.05). **(d)** Representative images of mitosis duration evaluated by live cell imaging using the Incucyte Zoom System in 22Rv1 cells after BUB1B depletion (vs control shGFP). Graphs show quantification of one representative experiment (50 randomly chosen cells per group), of two independent experiments (unpaired t-test, * p < 0.05).

### 3. BUB1B promotes *in vitro* and *in vivo* castration-resistant growth of androgen-dependent cells

To model castration resistant levels of BUB1B found in CRPC cells, BUB1B was stably expressed in two distinct ADPC cell lines, LNCaP and VCaP (Figure 3 a and Supplementary Figure 2), and the ability of BUB1B to promote proliferation in the absence of androgens (castration conditions) was evaluated. In both ADPC cell lines, ectopic expression of BUB1B was sufficient for cell growth under castration conditions whereas as expected, ADPC cells transduced with empty vector (EV) showed no or minimal growth (Figure 3 b). Moreover, ectopic BUB1B expression significantly increased anchorage-independent growth in soft agar assays under androgen-depleted conditions (Figure 3 c).

**Figure 3.**
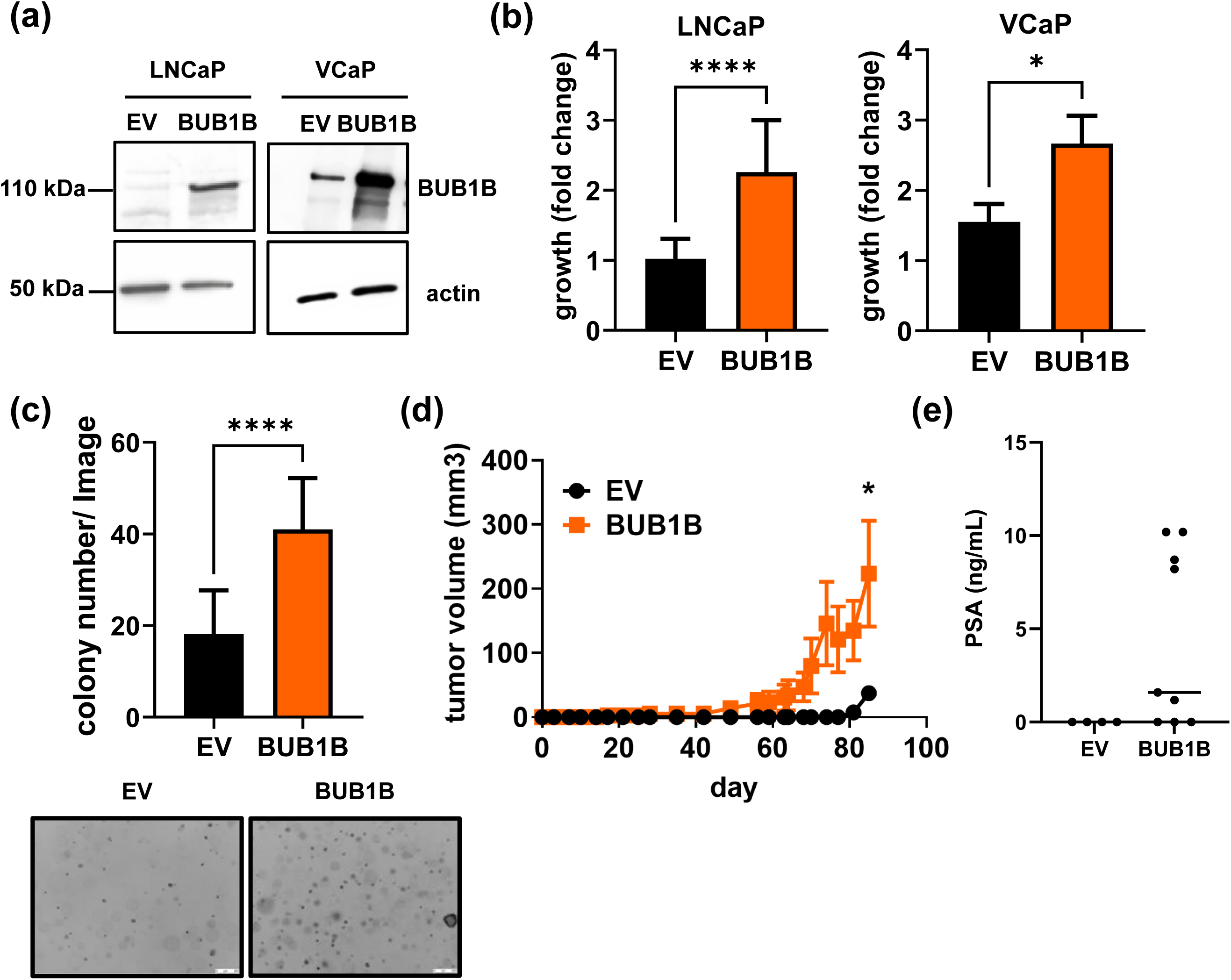
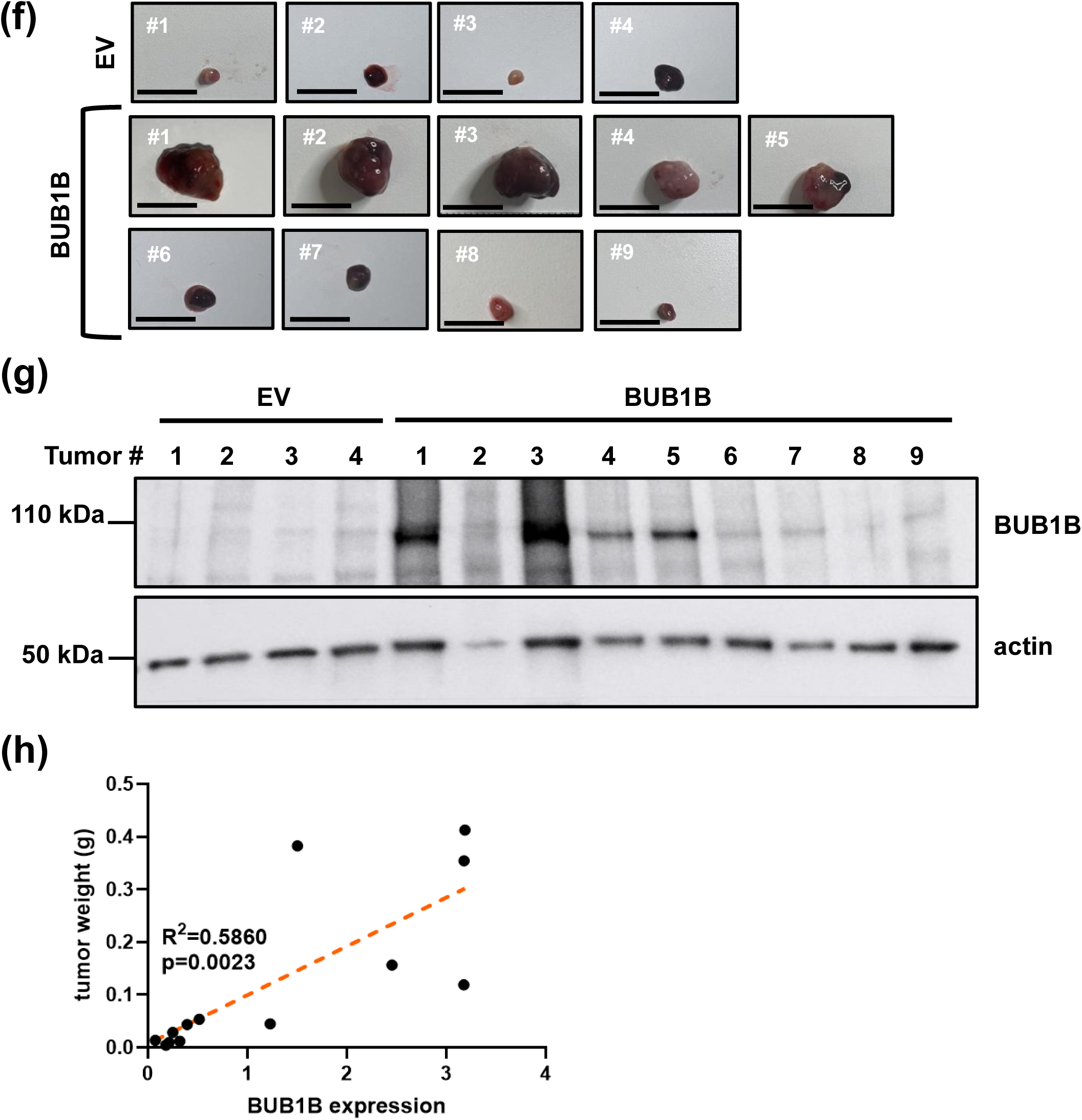
BUB1B ectopic expression in androgen-dependent PC cells is sufficient for castration-resistant growth. **(a)** BUB1B levels were determined by western blot of lysates from LNCaP and VCaP ADPC cells ectopically expressing BUB1B or EV. **(b)** LNCaP and VCaP BUB1B vs EV cell proliferation was measured by trypan blue exclusion in the absence of androgens after eight days. Graphs represent the average ± SEM of three or four independent experiments performed in triplicate (Mann-Whitney test, * p < 0.05). **(c)** Colony formation was evaluated in LNCaP EV and BUB1B in soft agar assays after fifteen days. Images are representative of one of three independent experiments. Graphs represent the average of three independent experiments performed in triplicate (unpaired t-test, **** p < 0.0001). **(d)** LNCaP EV or BUB1B expressing cells were subcutaneously xenografted in castrated SCID mice (n=10 for EV and n=29 for BUB1B) and tumor volumes (mm^3^) were measured by caliper every 3 days for 85 days. Tumor volume over time was evaluated using a mixed model with cubic effect. Differences were observed between groups at the end of the experiment (day 85). **(e)** End point PSA in plasma was determined by ELISA (Mann-Whitney test, **** p < 0.0001) **(f)** Images of end-stage tumors from LNCaP EV and BUB1B xenografts (scale bar = 1 cm). **(g)** BUB1B was evaluated by western blot in tumor tissue extracts. Tumor images and the corresponding tumor samples were ordered based on tumor size. **(h)** Graph shows the correlation between tumor weight and BUB1B expression (Pearson correlation, ** p < 0.01).

LNCaP cells stably expressing BUB1B or the EV control were xenografted into castrated SCID mice and tumor formation was evaluated. LNCaP BUB1B cells formed more rapidly growing tumors than EV cells. Furthermore, BUB1B tumors were significantly larger than EV cell tumors at end point (Figure 3 d and f). In accordance with larger tumor size, LNCaP BUB1B tumors tended to produce greater amounts of circulating PSA compared to EV tumors (Figure 3 e). Additionally, tumor weight correlated with BUB1B levels (Figure 3 g and h). Thus, BUB1B expression in ADPC cells was sufficient to promote castration resistant growth *in vitro* and in *vivo*.

### 4. BUB1B kinase activity is required for castration resistant growth

BUB1B WT or two previously characterized kinase-dead mutants (D882N and K795R) were ectopically expressed in LNCaP and VCaP ADPC cells to test whether BUB1B kinase activity was required for BUB1B-mediated progression to CRPC. Only BUB1B WT but not the BUB1B kinase dead mutants promoted growth in androgen-depleted conditions, indicating that BUB1B kinase activity was essential (Figure 4 a). Similarly, LNCaP BUB1B WT cells formed more colonies in soft agar than EV or the kinase dead mutant-expressing cells (Figure 4 b). Since BUB1B kinase activity was needed to promote ADPC progression to CRPC (Figure 4 a and b) and BUB1B depletion decreased CRPC proliferation (Figure 2), we performed ‘rescue’ experiments to determine whether BUB1B kinase activity was important for growth of established CRPC cells. Comparable levels of BUB1B WT and mutants were first expressed in 22Rv1 cells (Supplementary Figure 3 a). Growth of BUB1B WT and EV CRPC cells was the same indicating that BUB1B does not have general growth promoting effects. Furthermore, growth of CRPC cells ectopically expressing BUB1B WT and BUB1B kinase dead mutants was also comparable (Supplementary Figure 3 b), suggesting that kinase dead mutants did not have a dominant negative effect. Next, endogenous BUB1B levels were depleted (with siBUB1B against non-coding sequences) in WT, D882N, K795R or EV CRPC cells (Figure 4 c and d). Only the expression of BUB1B WT rescued the anti-proliferative effects of BUB1B depletion in 22Rv1 CRPC cells (Figure 4 c). These results show that BUB1B kinase activity was sufficient for ADPC progression to CRPC but also necessary to maintain growth of established CRPC cells.

**Figure 4.**
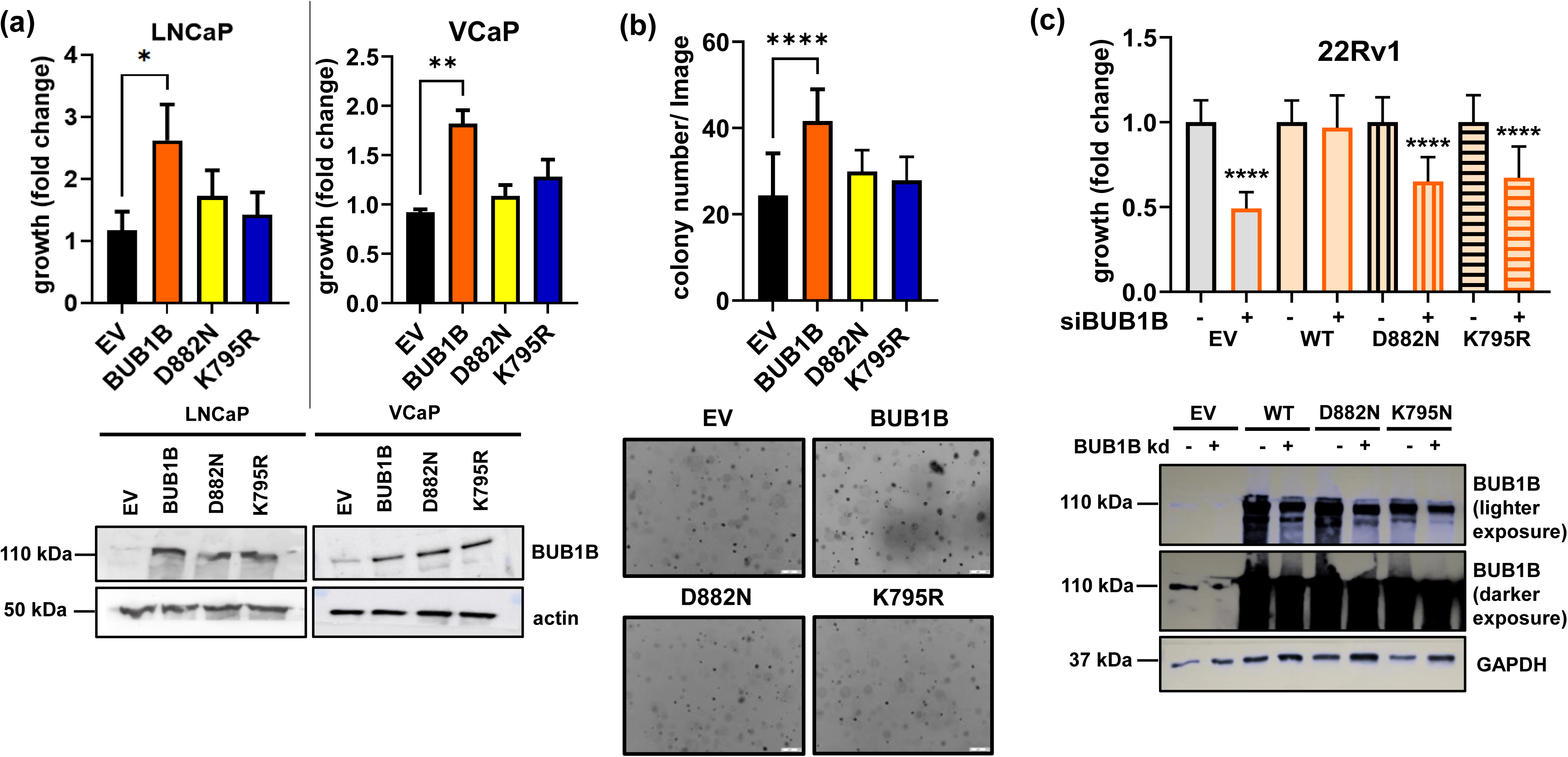
BUB1B kinase activity is required for BUB1B-mediated progression to CRPC and maintenance of CRPC. **(a)** LNCaP and VCaP ADPC cells expressing BUB1B (WT), D882N, K795R (kinase dead mutants) or control cells (EV) were grown in the absence of androgen and BUB1B protein was evaluated by western blot. Growth of LNCaP and VCaP expressing EV, BUB1B (WT), D882N and K795R was determined by trypan blue after eight days in culture. Graphs represent the average of three independent experiments performed in triplicate. (Kruskal-Wallis test, * p < 0.05, ** p < 0.01). **(b)** Colony formation of LNCaP EV, BUB1B, D882N and K795R was evaluated by soft agar assays after 3 weeks. Images are representative of three independent experiments (one-way ANOVA, * p < 0.05, **** p < 0.0001). BUB1B, D882N,K795R or EV control cells were expressed in 22Rv1 CRPC. **(c)** 22Rv1 cell proliferation was measured by trypan blue exclusion assay after BUB1B knockdown (siRNA against non-coding sequences) and ectopic expression of BUB1B, D882N,K795R or EV control. The graph represents the average of four independent experiments performed in triplicate (unpaired t-test, **** p < 0.0001). The lower panel shows BUB1B protein by western blotting in 22Rv1 EV, BUB1B, D882N and K795R expressing cells after knockdown of endogenous BUB1B using siBUB1B RNA targeting the 3’UTR.

### 5. BUB1B promotes castration resistant growth through an AR independent mechanism

Because AR pathway reactivation is a mechanism for CRPC progression, we addressed whether ectopic expression of BUB1B affected AR levels and/or activity. There were no appreciable differences in AR protein levels in LNCaP or VCaP BUB1B cells compared to EV cells (Figure 5 a and Supplementary Figure 4 a). In addition, depletion of AR using AR siRNA in LNCaP and VCaP EV and in their BUB1B counterparts showed that only EV, but not BUB1B cells, were growth inhibited (Figure 5 b Supplementary Figure 4 b). These data support a mechanism in which BUB1B promotion of castration resistant growth occurred by bypassing AR.

**Figure 5.**
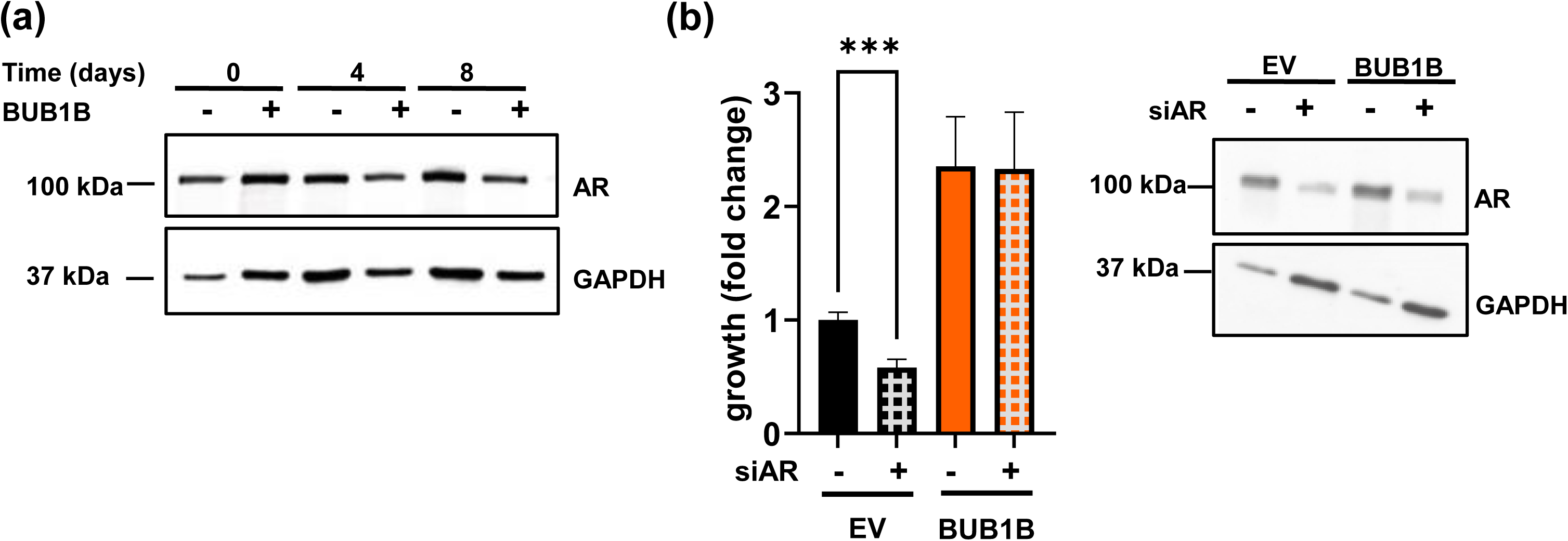
Castration-resistant growth of BUB1B cells does not require AR. **(a)** AR was evaluated by western blot in LNCaP EV and BUB1B cells growing in castrated conditions (2% CSS) at 0, 4 and 8 days. **(b)** AR was knocked down in LNCaP EV or BUB1B cells by siRNA and cell number was quantified by trypan blue exclusion assay after eight days. Graph represents three independent experiments performed in triplicate (Mann-Whitney test, * p < 0.05). AR levels after AR knockdown using siRNA in LNCaP EV and BUB1B cells are shown.

### 6. BUB1B expression promotes enzalutamide resistance

Because BUB1B-driven CRPC progression occurred in an AR independent manner, we evaluated whether BUB1B cells would be resistant to the AR antagonist, enzalutamide. Growth of LNCaP BUB1B and VCaP BUB1B cells was unaffected by enzalutamide, indicating that BUB1B promoted both castration- and enzalutamide resistance (Figure 6 a). Similarly, enzalutamide did not reduce the capacity of LNCaP BUB1B cells to form colonies in soft agar (Figure 6 b).

**Figure 6.**
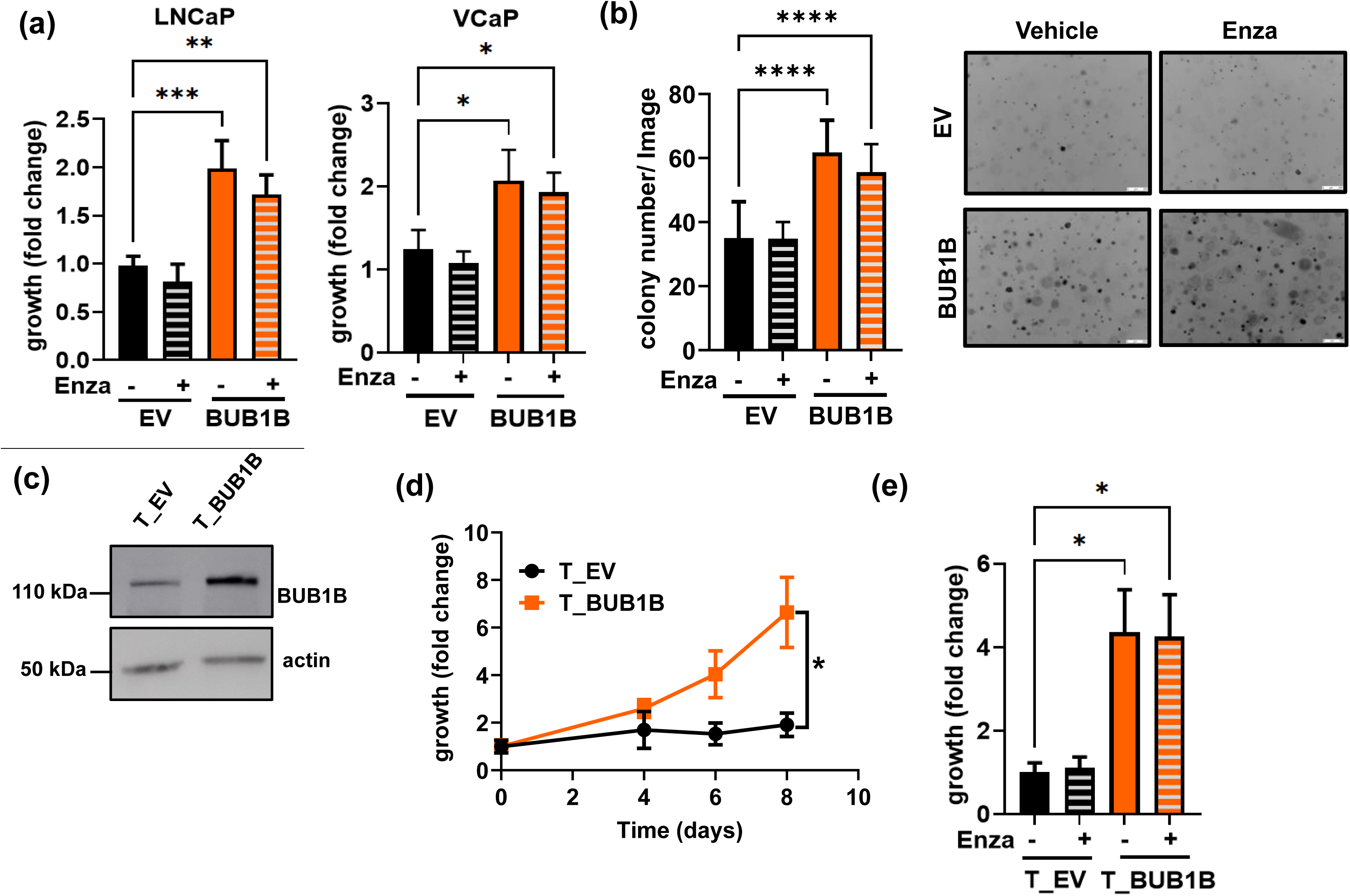
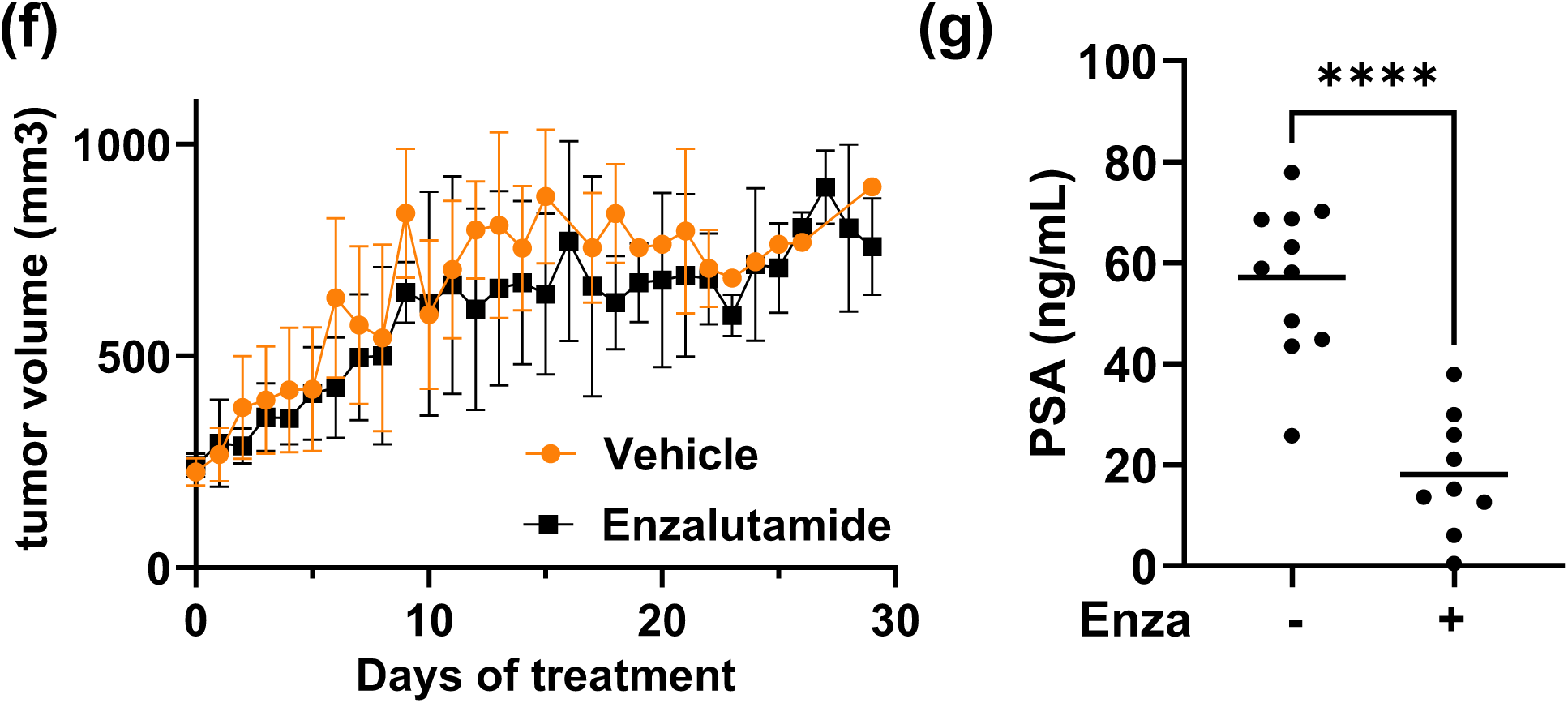
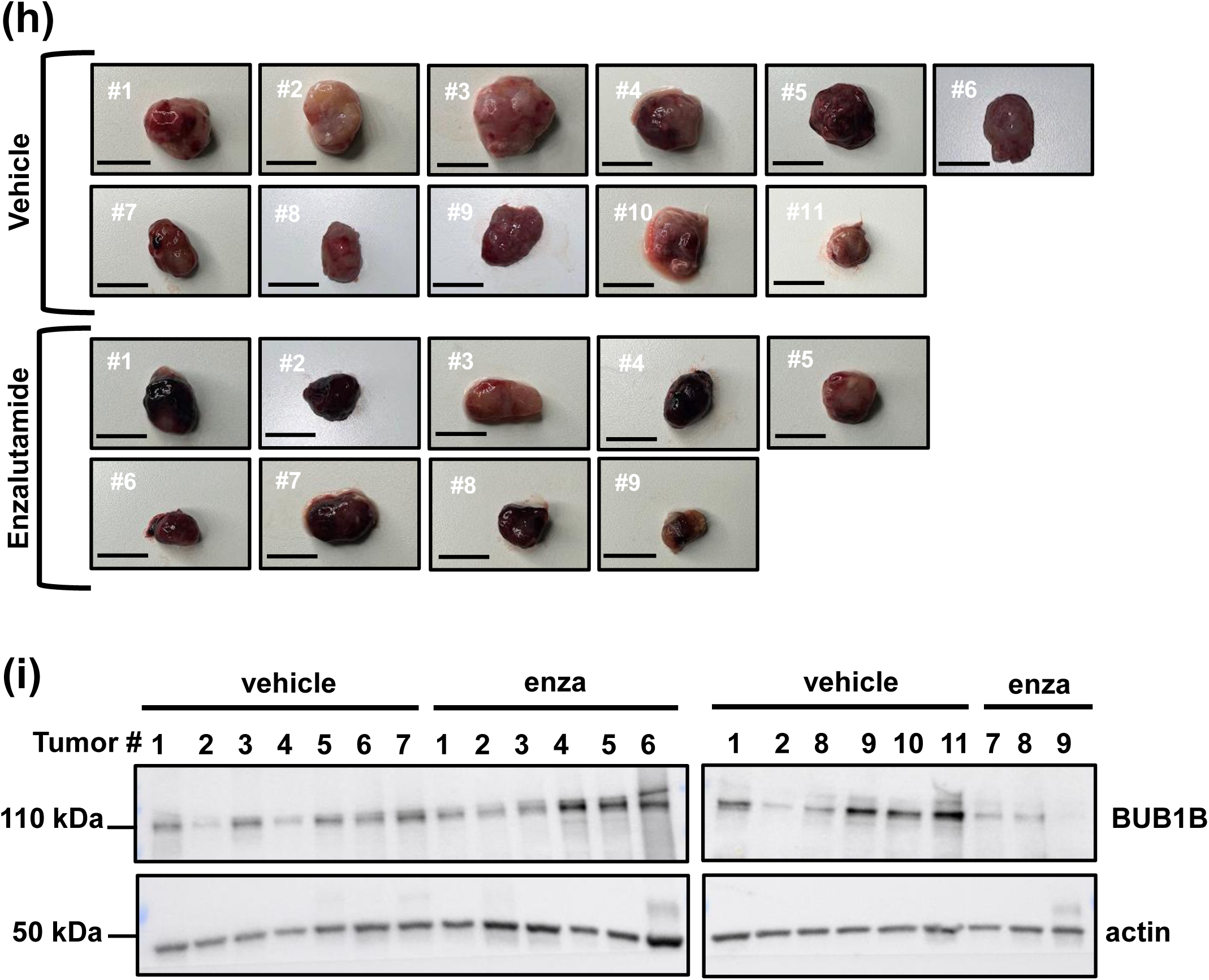
BUB1B drives enzalutamide resistance *in vitro* and *in vivo*. **(a)** Cell proliferation determined by trypan blue exclusion after eight days of enzalutamide treatment (10 µM) in LNCaP and VCaP EV and BUB1B cells. The graph represents three independent experiments done in triplicate (Mann-Whitney test, * p < 0.05). **(b)** Colony formation in soft agar was measured after fifteen days. Images are representative of three independent experiments performed in triplicate. Graphs are representative of three to four independent experiments performed in triplicate (one-way ANOVA, ** p < 0.01, **** p < 0.0001). **(c)** BUB1B was evaluated by western blot in LNCaP xenograft derived T_EV and T_BUB1B cells. **(d)** T_EV and T_BUB1B cells were grown in the absence of androgens and proliferation was measured by trypan blue exclusion assays. The graph represents three independent experiments performed in triplicates. The area under the curve was calculated (unpaired t-test, * p < 0.05). **(e)** Proliferation of T_EV and T_BUB1B cells treated with 10 µM enzalutamide (or vehicle) for eight days is shown. The graph represents three independent experiments performed in triplicate (Kruskal-Wallis test, * p < 0.05). **(f)** LNCaP EV or T_BUB1B expressing cells were subcutaneously xenografted into castrated SCID mice (EV n=10 and T_BUB1B n=29) and tumor volumes (mm3) were measured by caliper every 3 days. Once tumors reach 200 mm3, animals were split into two groups and treated daily with enzalutamide (n=9) or vehicle (n=11). Graph shows tumor volume over time. **(g)** PSA levels in plasma were determined by ELISA (Mann-Whitney test, **** p < 0.0001). **(h)** Images of end-stage tumors from T_BUB1B vehicle (n=11) or enzalutamide (n=9) xenografts are shown. **(i)** BUB1B protein was evaluated in tumors by western blot. Tumor images and protein samples were ordered based on tumor size, vehicle tumors #1 and 2 were repeated in both blots for comparison.

We generated and authenticated LNCaP cell lines (termed T_EV and T_BUB1B) from end-stage castration resistant tumors shown in figure 3. The xenograft tumor-derived T_BUB1B cells retained higher BUB1B expression compared to T_EV controls (Figure 6 c). Furthermore, only T_BUB1B cells were able to proliferate following enzalutamide treatment *in vitro* (Figure 6 d and e).

Since LNCaP BUB1B and VCaP BUB1B cells grew in the presence of enzalutamide (Figure 6), we evaluated expression of AR-V7 and GR, two well described mechanisms of enzalutamide resistance ^32,33^. LNCaP BUB1B cells, like their EV counterparts, remained AR-V7 and GR negative (Supplementary Figure 5 a). Similarly, VCaP BUB1B cells did not show changes in AR-V7 or GR levels compared to VCaP EV (Supplementary Figure 5 b). Moreover, treatment with the GR antagonist, RU486 (mifepristone), did not inhibit LNCaP BUB1B cell growth whereas PC3 cells, which express endogenous GR, were growth inhibited by RU486 (Supplementary Figure 5 c). These results suggest that neither AR-V7 nor GR participate in BUB1B-dependent CRPC growth and enzalutamide resistance. Neuroendocrine differentiation is another pathway leading to enzalutamide resistance; however, levels of synaptophysin, a well-described NEPC marker, were comparatively low and unchanged by BUB1B expression in LNCaP and VCaP BUB1B and in T_BUB1B cells compared to NCI-H660 an established NEPC cell model (Supplementary Figure 6). To evaluate the effect of enzalutamide *in vivo*, T_BUB1B cells or the empty vector (EV) controls were xenografted into castrated SCID mice. T_BUB1B cells formed tumors with ∼90% tumor take, whereas EV cells resulted in only one small, breakthrough tumor out of 10 possible tumor sites (Supplementary Figure 7 a). Once T_BUB1B tumors reached 200 mm^3^, animals were divided into two groups and treated daily with enzalutamide (n=9 animals) or vehicle (n=11 animals). T_BUB1B tumors did not significantly respond to treatment, evidenced by no differences in tumor volume after 30 days of enzalutamide treatment (Figure 6 f and Supplementary Figure 7 b). Circulating PSA levels were significantly reduced in the enzalutamide treated group (Figure 6 g) consistent with enzalutamide-mediated inhibition of AR suggesting an uncoupling of PSA and tumor growth. Figure 6 i shows images of the T_BUB1B xenografts at the experimental end point confirming no major differences in tumor sizes between mice treated with enzalutamide or vehicle. While BUB1B levels across all tumor samples were variable, there were no obvious differences between the enzalutamide-treated compared to vehicle control groups (Figure 6 i). Taken together, our results show that BUB1B expression promoted CRPC progression and enzalutamide resistance in vitro and in vivo.

### 7. Inhibition of the BUB1B substrate CENP-E blocks CRPC and enzalutamide resistant growth

The kinesin-like mitotic protein CENP-E is the only known BUB1B phospho-substrate (aside from BUB1B itself). To interrogate further the role of BUB1B kinase activity in CRPC progression, we performed proof-of-concept growth assays using the CENP-E inhibitor GSK-923295 as there are no available BUB1B inhibitors. We found that growth of LNCaP and VCaP BUB1B and LNCaP T_BUB1B cell lines were decreased following CENP-E inhibition whereas, as expected, there was minimal effect on the non-proliferative EV cells (Figure 7 a and b, Supplementary Figure 8 a). These results further support the importance of BUB1B kinase activity in CRPC growth. Next BUB1B cells were treated with different doses of GSK-923295 and enzalutamide individually or in combination. Dual treatment significantly decreased viability of BUB1B cells (Supplementary Figure 8 b). Moreover, studies with combinations of different doses of both inhibitors showed that BUB1B cells were synergistically inhibited by GSK-923295 and enzalutamide (combinatory index, CI < 1) (Figure 7 c). In 22Rv1 CRPC cells, which are relatively resistant to enzalutamide, GSK-923295 increased sensitivity to enzalutamide resulting in significantly reduced proliferation (Supplementary Figure 8 c). These results support the importance of BUB1B kinase / CENP-E axis in castration- and enzalutamide-resistance.

**Figure 7.**
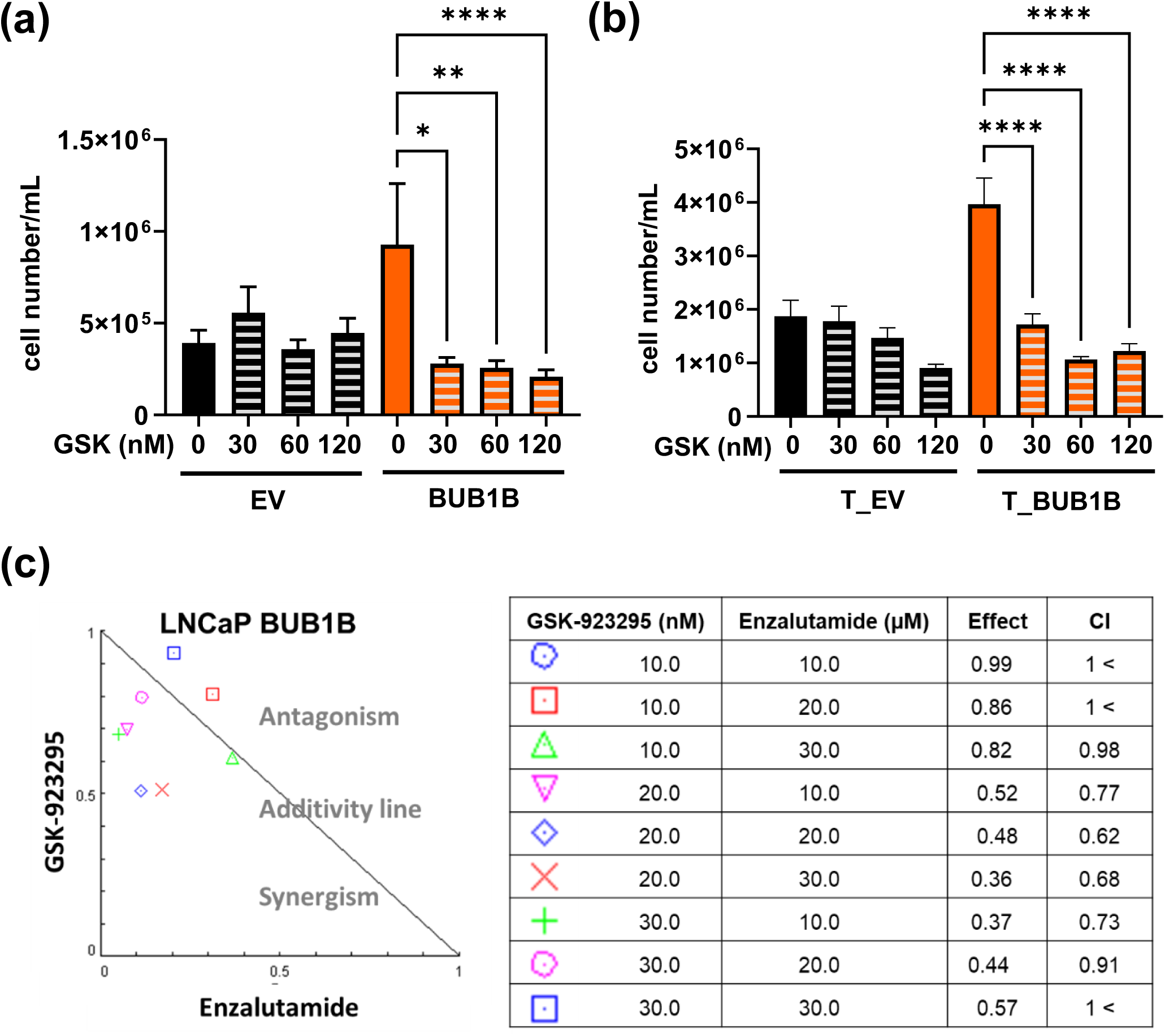
Inhibition of BUB1B downstream target CENP-E blocks progression to CRPC and enzalutamide resistance. Cell proliferation was measured after GSK-923295 (CENP-E inhibitor) treatment (0, 0.03, 0.06, 0.12 and 0.25 µM) for 7 days in **(a)** LNCaP EV and BUB1B cells or **(b)** LNCaP T_EV and T_BUB1B cells. Graph represents four independent experiments performed in triplicate (one-way ANOVA, * p < 0.05, ** p < 0.01). **(c)** BUB1B cells growing in castration conditions were treated with GSK-923295 (10-30 nM), enzalutamide (10-30 µM) or the combination for 7 days and cell number was evaluated by trypan blue exclusion assay. Two independent experiments were performed in triplicate. Combinatory index (CI) was calculated using the CompuSyn software. A CI < 1 indicates synergism, = 1 indicates additivity,> 1 indicates antagonism.

## Discussion

New CRPC therapeutic options are needed, particularly for men whose disease progresses despite maximal AR blockade with drugs such as enzalutamide. We identified roles for BUB1B kinase in both progression to enzalutamide-resistant CRPC and in maintenance of the CRPC state, supporting BUB1B as a promising therapeutic target. Ectopic BUB1B expression in ADPC cells, at levels observed in CRPC, was sufficient to promote CRPC progression and enzalutamide resistance, whereas BUB1B depletion halted CRPC proliferation through cell cycle arrest and mitosis delay. Mechanistically, we demonstrated that BUB1B kinase activity was required for in vitro and in vivo progression to enzalutamide-resistant CRPC and CRPC growth. Because BUB1B is more highly expressed in advanced PC and that CRPC cell proliferation was dependent on BUB1B kinase activity, BUB1B inhibition is a promising approach for CRPC and enzalutamide-resistant disease.

BUB1B was higher in PC and CRPC patient samples (publicly available RNA-seq data and in-house tumor microarray analysis), and this was reflected in PC and CRPC cell lines as well as castration-resistant PDXs. Higher BUB1B expression was also associated with worse prognosis in patients (Figure 1). We demonstrated that BUB1B depletion inhibited CRPC proliferation to a greater extent than in ADPC cells, suggesting an association between advanced disease and dependency on BUB1B. Thus, there may be a therapeutic window for BUB1B inhibition in treatment-resistant, aggressive CRPC. Increased levels of mitotic errors and aneuploidy are correlated with progression and aggressiveness of CRPC^34,35^. In this context, it has been proposed that cancer cells develop mechanisms to tightly regulate mitosis (such as by increased expression of mitotic kinases) and maintain mitotic errors within a tolerable range^36^. Mitotic regulators therefore represent a potential vulnerability for aggressive tumors^37–39^. In agreement with these concepts, we showed that BUB1B knockdown arrested CRPC cells in G2/M phase and prolonged mitosis (Figure 2), suggesting that BUB1B promotes CRPC proliferation through its canonical function in mitosis regulation. Similarly in breast cancer cells, BUB1B knockdown decreased cell proliferation through cell cycle arrest *in vitro* and *in vivo*^40,41^. Mechanistically, rescue experiments performed in established CRPC cells indicated that BUB1B kinase activity was required to sustain proliferation (Figure 6); further supporting the requirement of tighter mitosis regulation in CRPC. BUB1B may drive CRPC through increased kinase activity which would serve to strengthen the mitotic checkpoint and restrict segregation errors during progression.

Enzalutamide resistance occurs through either AR-dependent or AR-independent mechanisms^10,42–44^. While increased AR or AR-Vs expression can drive enzalutamide resistance, neither full length AR nor AR-V7 protein were elevated in BUB1B expressing PC cells (Figure 5 and Supplementary Figure 4 and 5). Importantly, BUB1B-mediated progression to CRPC was unaffected by AR depletion (Figure 5). Also, in line with an AR independent mechanism of enzalutamide resistance, BUB1B depletion decreased CRPC cell proliferation irrespective of AR and AR-V7 status (Figure 2). These results support BUB1B promotion of aggressive PC growth as a pervasive mechanism. Strong pressure exerted by treatment with AR signaling inhibitors (e.g enzalutamide) can result in GR upregulation and hijacking of the oncogenic AR transcriptional program by GR. However, BUB1B expression did not induce GR levels nor were LNCaP BUB1B cells sensitive to the potent GR/progesterone receptor antagonist, RU486 (Supplementary Figure 4). Similarly, under maximal AR blockade, lineage plasticity is increasingly implicated in promotion of neuroendocrine differentiation concomitant with decreased AR dependence. Neither did we observe neuroendocrine differentiation as assessed by synaptophysin (Supplementary Figure 5) nor were AR levels reduced in BUB1B cells (Figure 5), suggesting that neuroendocrine differentiation was not involved in BUB1B-dependent enzalutamide resistance.

The dependency on BUB1B kinase activity for CRPC proliferation poses kinase inhibition as a tractable approach. Currently, there are no commercially available BUB1B kinase inhibitors, however recent *in silico* studies identified bioactive compounds predicted to bind BUB1B’s kinase domain and these may be useful in designing BUB1B specific inhibitors47. Other potential BUB1B kinase inhibitors have been synthesized and shown to inhibit BUB1B enzymatic activity in biochemical assays48. Similarly, bubristatin was identified as a small-molecule tool to target BUB1B kinase domain17. However, these BUB1B inhibitors have not been used for animal studies.

We showed that BUB1B with functional kinase activity (and not kinase deficient mutants) was sufficient for progression to CRPC and for maintaining CRPC growth (Figure 4). In line with these findings, we found that inhibition of CENP-E, the only known BUB1B substrate (aside from itself), decreased CRPC proliferation and sensitized BUB1B cells to enzalutamide (Figure 7). Impairment of BUB1B kinase activity is known to cause aberrant CENP-E localization and chromosome segregation errors during mitosis^16,18,19^. Moreover, CENP-E inhibition (using siRNA or GSK-923295) blocks CRPC growth *in vitro* and *in vivo*^45^. That BUB1B kinase activity is a CRPC vulnerability is consistent with emerging data demonstrating that mitotic regulators, including BUB1B, are associated with progression to and aggressiveness of CRPC^36^. Recent reports described that inhibition of AR pathway using abiraterone induces mild mitotic defects^46^. Similarly, enzalutamide treatment prolongs mitosis duration in PC cell lines^47^. Since BUB1B kinase activity was required to promote growth in ADT conditions (Figure 4) and it was also required for enzalutamide resistance development (Figure 7); BUB1B phosphorylation of CENP-E may mediate resistance to enzalutamide by correcting treatment-induced mitotic defects. Then, disruption of the BUB1B/ CENP-E axis may represent a therapeutic alternative for enzalutamide resistant CRPC.

In summary, our findings demonstrate that the BUB1B kinase pathway is an untapped susceptibility that is a potential therapeutic strategy for advanced and enzalutamide-resistant CRPC.

## Acknowledgments

We are grateful to Dr. Colm Morrissey, PhD (Department of Urology, University of Washington) for providing the TMAs containing PC metastases. We thank Morgan Pantone for her input during the manuscript preparation. We thank Dr. Wei Zhao from Sylvester Biostatistics and Bioinformatics Shared Resource (BBSR) for performing the statistical analysis of the *in vivo* work. This work was supported by Sylvester Comprehensive Cancer Center Tumor Biology Research Grant for Trainees FY2020, awarded to Maria Julia Martinez. This work was supported by VA Merit Review 2I01BX002773-06 from the United States Department of Veterans Affairs, Biomedical Laboratory Research and Development Service, awarded to Kerry L Burnstein, PhD, Research Health Scientist, Bruce W. Carter Department of Veterans Affairs Medical Center, Miami, FL.

## Author contributions

M.J.M. was responsible for experimental design, conducting research, data analysis, and writing. C.B, N.P. and R.D.Z.L. conducted research and data analysis. M.V.R conducted data analysis. K.L.B. was responsible for experimental design, data analysis, providing funding, and writing.

## Supplementary Figure legends

**<u>Supplementary Figure 1. BUB1B knock down decreases CRPC proliferation.</u> (a)** 22Rv1 cell proliferation was measured by trypan blue exclusion assay after BUB1B depletion using three different shRNA constructs [two against the 3’ UTR and one against the coding (CDR) regions]. Graph represents one independent experiment performed in quadruplicate. BUB1B in 22Rv1 cells transduced with shGFP control and 3’ UTR#1, 3’UTR #2 and CDR was measured by Western blot. **(b)** Graphs represent the quantification of western blots (shown in Figure 5b) for cyclin A and CDK1 in 22Rv1 after BUB1B depletion. **(c)** CDK1 levels were evaluated in LNCaP and C4-2B BUB1B depleted cells by Western blot. cl-PARP levels were evaluated in 22Rv1 BUB1B depleted cells after **(d)** one, two or three days and **(e)** after seven days. **(f)** Caspase -3/ -7 activity was evaluated in 22Rv1 cells after BUB1B depletion. Docetaxel was used as a positive control. Graphs represent one of two independent experiments performed in eight technical replicates.

**<u>Supplementary Figure 2. Ectopically expressed BUB1B in LNCaP and VCaP ADPC</u> <u>cells is comparable to CRPC cells.</u>** BUB1B was expressed in LNCaP and VCaP cells, and BUB1B levels were compared to 22Rv1 and C42B CRPC cells by western blot.

**<u>Supplementary Figure 3. Ectopically expressed BUB1B (WT or kinase dead</u> <u>mutants) does not promote growth of 22Rv1 CRPC cells</u>. (a)** BUB1B WT, D882N, K795R protein levels were measured by Western blot in 22Rv1 cells. **(b)** Cell proliferation was evaluated by trypan blue exclusion assay in 22Rv1 expressing EV, BUB1B, D882N and K795R. Graphs represent three to four independent experiments performed in triplicate.

**<u>Supplementary Figure 4. Ectopically expressed BUB1B in VCaP ADPC cells</u> <u>promotes castration-resistant growth through AR bypass.</u> (a)** AR protein was evaluated by western blot in VCaP EV and BUB1B growing in castration conditions. **(b)** AR was depleted using siRNA in VCaP EV and BUB1B. Cells were cultured in castration conditions and growth was evaluated after 8 days by trypan blue exclusion. Graph represents one independent experiment performed in triplicate. AR knockdown efficiency was evaluated by western blot.

**<u>Supplementary Figure 5. BUB1B promotes enzalutamide resistance independent</u> <u>of AR-V7 or GR.</u> (a)** AR-V7 and GR protein were evaluated in LNCaP EV and BUB1B cells after enzalutamide treatment at different time points (0, 4, 8 days). 22Rv1 cells were used as control for AR-V7 and GR expression. **(b)** GR, AR and AR-Vs were determined by western blot in VCaP EV and BUB1B growing in castration conditions after 8 days**.(c)** Cells were treated with the GR inhibitor mifepristone (5 µM) for 8 days in androgen-depleted conditions. PC3 cells were used as positive control. Graph represents one independent experiment performed in triplicate.

**<u>Supplementary Figure 6. Ectopically expressed BUB1B does not result in</u> <u>neuroendocrine differentiation.</u>** Synaptophysin was evaluated by western blot in **(a)** LNCaP and VCaP EV and BUB1B cells and **(b)** T_EV and T_BUB1B cells. Graphs show the quantification of two to four independent experiments. NCI-H660 (a human NEPC cell line) was used as positive control.

**<u>Supplementary Figure 7. BUB1B drives enzalutamide resistance *in vivo*.</u>** Spaghetti plots show tumor volumes of **(a)** T_BUB1B over time (85 days) or **(b)** Mice bearing T_BUB1B xenografts were treated with vehicle (orange) or enzalutamide (black).

**<u>Supplementary Figure 8. CENP-E inhibition sensitizes BUB1B cells to</u> <u>enzalutamide.</u> (a)** VCaP EV and BUB1B cells were treated with GSK-923295 (20 nM) and cell proliferation was evaluated after 7 days of treatment. Graph represents three independent experiments performed in triplicate. **(b)** LNCaP EV and BUB1B cells growing in castration conditions were treated with GSK-923295 (10-30 nM), enzalutamide (10-30 µM) or their combination for 7 days and cell number was evaluated by trypan blue exclusion assay. Graphs represent two independent experiments performed in triplicate (one-way ANOVA, * p < 0.05, ** p < 0.01, *** p < 0.001). **(c)** 22Rv1 cells were treated with GSK-923295 (20 nM), enzalutamide (20 µM) individually or in combination. Graph represents three independent experiments performed in triplicate (one-way ANOVA, * p < 0.05).

## References

1. Abida, W., et al. Genomic correlates of clinical outcome in advanced prostate cancer. Proc Natl Acad Sci U S A 116, 11428–11436 (2019).

2. Dreicer, R. Managing advanced prostate cancer: the rapidly changing treatment landscape. Am J Manag Care 20, S282–289 (2014).

3. Grasso, C.S., et al. The mutational landscape of lethal castration-resistant prostate cancer. Nature 487, 239–243 (2012).

4. Nouruzi, S., Kobelev, M., Tabrizian, N., Gleave, M. & Zoubeidi, A. New frontiers in prostate cancer treatment from systemic therapy to targeted therapy. EMBO Mol Med 17, 2191–2214 (2025).

5. Desai, K., McManus, J.M. & Sharifi, N. Hormonal Therapy for Prostate Cancer. Endocr Rev 42, 354–373 (2021).

6. Paschalis, A. & de Bono, J.S. Prostate Cancer 2020: “The Times They Are a’Changing“. Cancer Cell 38, 25–27 (2020).

7. Siegel, R.L., Miller, K.D., Wagle, N.S. & Jemal, A. Cancer statistics, 2023. CA Cancer J Clin 73, 17–48 (2023).

8. Tilki, D., Schaeffer, E.M. & Evans, C.P. Understanding Mechanisms of Resistance in Metastatic Castration-resistant Prostate Cancer: The Role of the Androgen Receptor. Eur Urol Focus 2, 499–505 (2016).

9. Crona, D.J. & Whang, Y.E. Androgen Receptor-Dependent and -Independent Mechanisms Involved in Prostate Cancer Therapy Resistance. Cancers (Basel) 9(2017).

10. Dai, C., Dehm, S.M. & Sharifi, N. Targeting the Androgen Signaling Axis in Prostate Cancer. J Clin Oncol 41, 4267–4278 (2023).

11. Magani, F., et al. Identification of an oncogenic network with prognostic and therapeutic value in prostate cancer. Mol Syst Biol 14, e8202 (2018).

12. Bolanos-Garcia, V.M. & Blundell, T.L. BUB1 and BUBR1: multifaceted kinases of the cell cycle. Trends Biochem Sci 36, 141–150 (2011).

13. Faesen, A.C., et al. Basis of catalytic assembly of the mitotic checkpoint complex. Nature 542, 498–502 (2017).

14. Musacchio, A. The Molecular Biology of Spindle Assembly Checkpoint Signaling Dynamics. Curr Biol 25, R1002–1018 (2015).

15. Suijkerbuijk, S.J., et al. The vertebrate mitotic checkpoint protein BUBR1 is an unusual pseudokinase. Dev Cell 22, 1321–1329 (2012).

16. Guo, Y., Kim, C., Ahmad, S., Zhang, J. & Mao, Y. CENP-E--dependent BubR1 autophosphorylation enhances chromosome alignment and the mitotic checkpoint. J Cell Biol 198, 205–217 (2012).

17. Huang, Y., et al. BubR1 phosphorylates CENP-E as a switch enabling the transition from lateral association to end-on capture of spindle microtubules. Cell Res 29, 562–578 (2019).

18. Weber, J., et al. A conserved CENP-E region mediates BubR1-independent recruitment to the outer corona at mitotic onset. Curr Biol 34, 1133–1141 e1134 (2024).

19. Ciossani, G., et al. The kinetochore proteins CENP-E and CENP-F directly and specifically interact with distinct BUB mitotic checkpoint Ser/Thr kinases. J Biol Chem 293, 10084–10101 (2018).

20. Alwadi, D., et al. RNA-Seq Uncovers Association of Endocrine-Disrupting Chemicals with Hub Genes and Transcription Factors in Aggressive Prostate Cancer. Int J Mol Sci 26(2025).

21. Modanwal, S., Mulpuru, V., Mishra, A. & Mishra, N. Transcriptomic signatures of prostate cancer progression: a comprehensive RNA-seq study. 3 Biotech 15, 135 (2025).

22. Liu, H., et al. Construction of Potential Gene Expression and Regulation Networks in Prostate Cancer Using Bioinformatics Tools. Oxid Med Cell Longev 2021, 8846951 (2021).

23. Cirak, Y., et al. Predictive and prognostic values of Tau and BubR1 protein in prostate cancer and their relationship to the Gleason score. Med Oncol 30, 526 (2013).

24. Fu, X., et al. Overexpression of BUB1B contributes to progression of prostate cancer and predicts poor outcome in patients with prostate cancer. Onco Targets Ther 9, 2211–2220 (2016).

25. Tian, J.H., et al. BUB1B Promotes Proliferation of Prostate Cancer via Transcriptional Regulation of MELK. Anticancer Agents Med Chem 20, 1140–1146 (2020).

26. Luca, B.A., et al. DESNT: A Poor Prognosis Category of Human Prostate Cancer. Eur Urol Focus 4, 842–850 (2018).

27. Cai, C., et al. ERG induces androgen receptor-mediated regulation of SOX9 in prostate cancer. J Clin Invest 123, 1109–1122 (2013).

28. Lyons, L.S., et al. Ligand-independent activation of androgen receptors by Rho GTPase signaling in prostate cancer. Mol Endocrinol 22, 597–608 (2008).

29. Magani, F., et al. Targeting AR Variant-Coactivator Interactions to Exploit Prostate Cancer Vulnerabilities. Mol Cancer Res 15, 1469–1480 (2017).

30. Zhao, N., et al. Arginine vasopressin receptor 1a is a therapeutic target for castration-resistant prostate cancer. Sci Transl Med 11(2019).

31. Nguyen, H.M., et al. LuCaP Prostate Cancer Patient-Derived Xenografts Reflect the Molecular Heterogeneity of Advanced Disease an--d Serve as Models for Evaluating Cancer Therapeutics. Prostate 77, 654–671 (2017).

32. Watson, P.A., Arora, V.K. & Sawyers, C.L. Emerging mechanisms of resistance to androgen receptor inhibitors in prostate cancer. Nat Rev Cancer 15, 701–711 (2015).

33. He, Y., et al. Androgen receptor splice variants bind to constitutively open chromatin and promote abiraterone-resistant growth of prostate cancer. Nucleic Acids Res 46, 1895–1911 (2018).

34. Stopsack, K.H., et al. Aneuploidy drives lethal progression in prostate cancer. Proc Natl Acad Sci U S A 116, 11390–11395 (2019).

35. Miller, E.T., et al. Chromosomal instability in untreated primary prostate cancer as an indicator of metastatic potential. BMC Cancer 20, 398 (2020).

36. Dhital, B., et al. Harnessing transcriptionally driven chromosomal instability adaptation to target therapy-refractory lethal prostate cancer. Cell Rep Med 4, 100937 (2023).

37. Cohen-Sharir, Y., et al. Aneuploidy renders cancer cells vulnerable to mitotic checkpoint inhibition. Nature 590, 486–491 (2021).

38. Quinton, R.J., et al. Whole-genome doubling confers unique genetic vulnerabilities on tumour cells. Nature 590, 492–497 (2021).

39. Serrano-Del Valle, A., et al. Future prospects for mitosis-targeted antitumor therapies. Biochem Pharmacol 190, 114655 (2021).

40. Koyuncu, D., Sharma, U., Goka, E.T. & Lippman, M.E. Spindle assembly checkpoint gene BUB1B is essential in breast cancer cell survival. Breast Cancer Res Treat 185, 331–341 (2021).

41. Lu, Y., et al. Downregulation of BUBR1 regulates the proliferation and cell cycle of breast cancer cells and increases the sensitivity of cells to cisplatin. In Vitro Cell Dev Biol Anim 59, 778–789 (2023).

42. Blatt, E.B. & Raj, G.V. Molecular mechanisms of enzalutamide resistance in prostate cancer. Cancer Drug Resist 2, 189–197 (2019).

43. Wang, Y., et al. Mechanisms of enzalutamide resistance in castration-resistant prostate cancer and therapeutic strategies to overcome it. Br J Pharmacol 178, 239–261 (2021).

44. Miller, C.D., Likasitwatanakul, P., Toye, E., Hwang, J.H. & Antonarakis, E.S. Current uses and resistance mechanisms of enzalutamide in prostate cancer treatment. Expert Rev Anticancer Ther 24, 1085–1100 (2024).

45. Liang, Y., et al. LSD1-Mediated Epigenetic Reprogramming Drives CENPE Expression and Prostate Cancer Progression. Cancer Res 77, 5479–5490 (2017).

46. Patterson, J.C., et al. Plk1 Inhibitors and Abiraterone Synergistically Disrupt Mitosis and Kill Cancer Cells of Disparate Origin Independently of Androgen Receptor Signaling. Cancer Res 83, 219–238 (2023).

47. Williams, C.S., et al. Inhibition of Androgen Receptor Exposes Replication Stress Vulnerability in Prostate Cancer. bioRxiv (2024).

