## Supplementary Figures for "BUB1B MITOTIC KINASE DRIVES THERAPY RESISTANT PROSTATE CANCER"

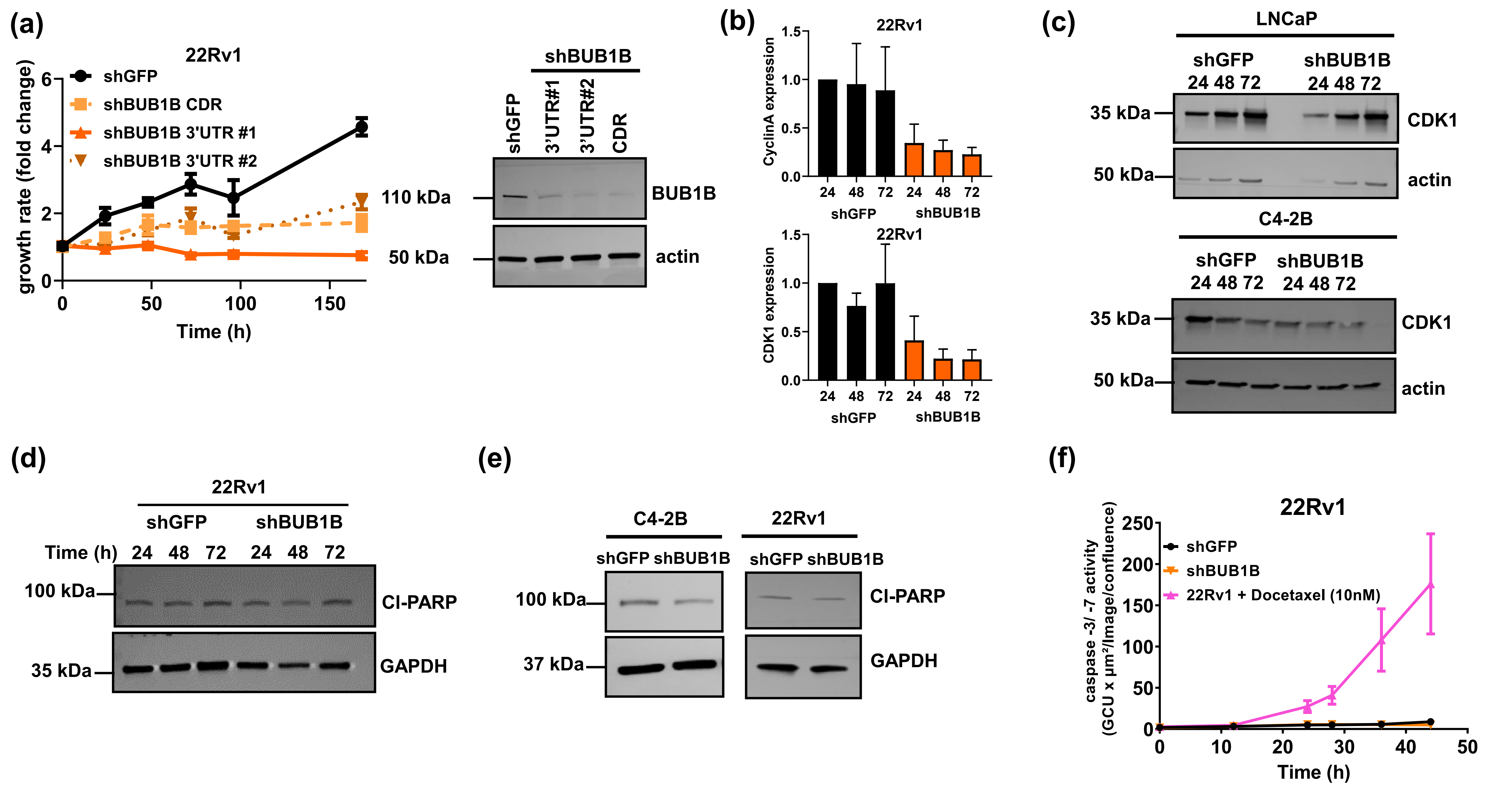


### Supplementary Figure 1

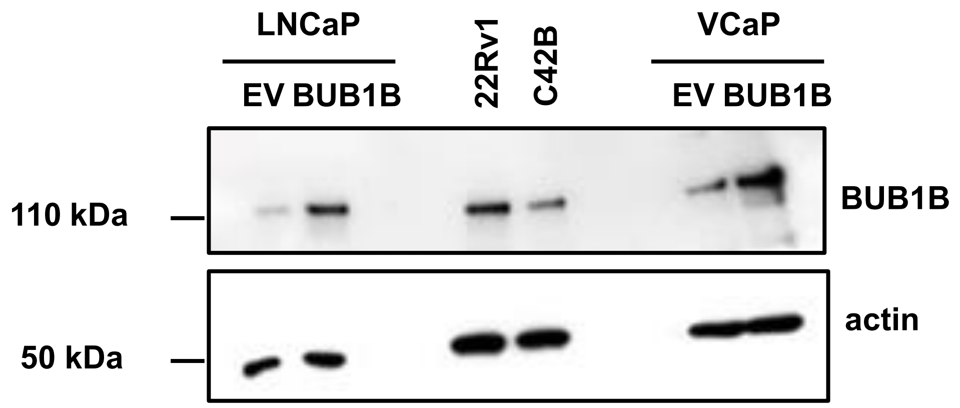


**Supplementary Figure 2**

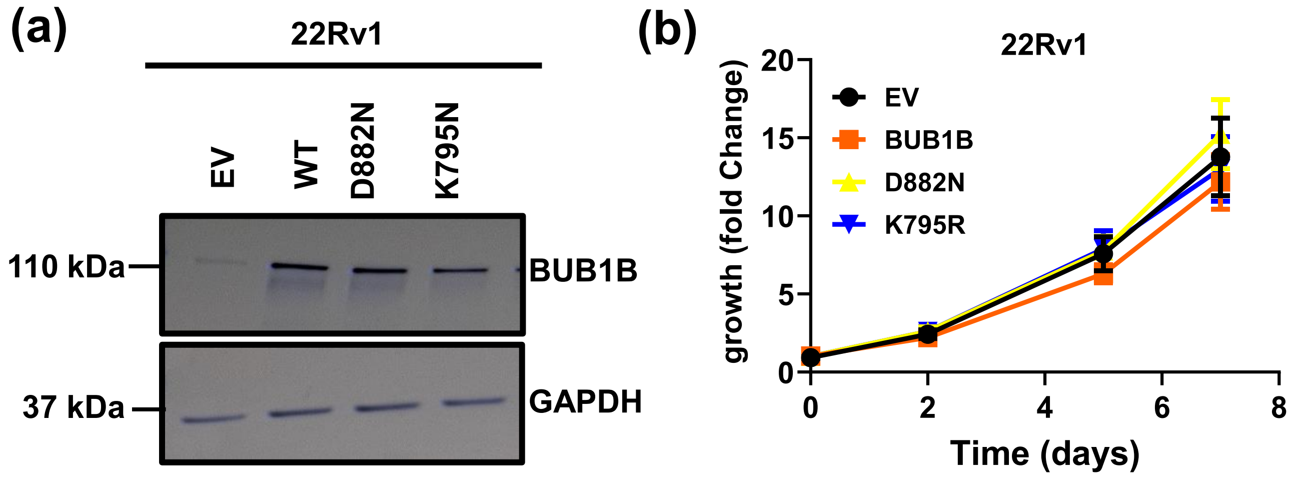


**Supplementary Figure 3**

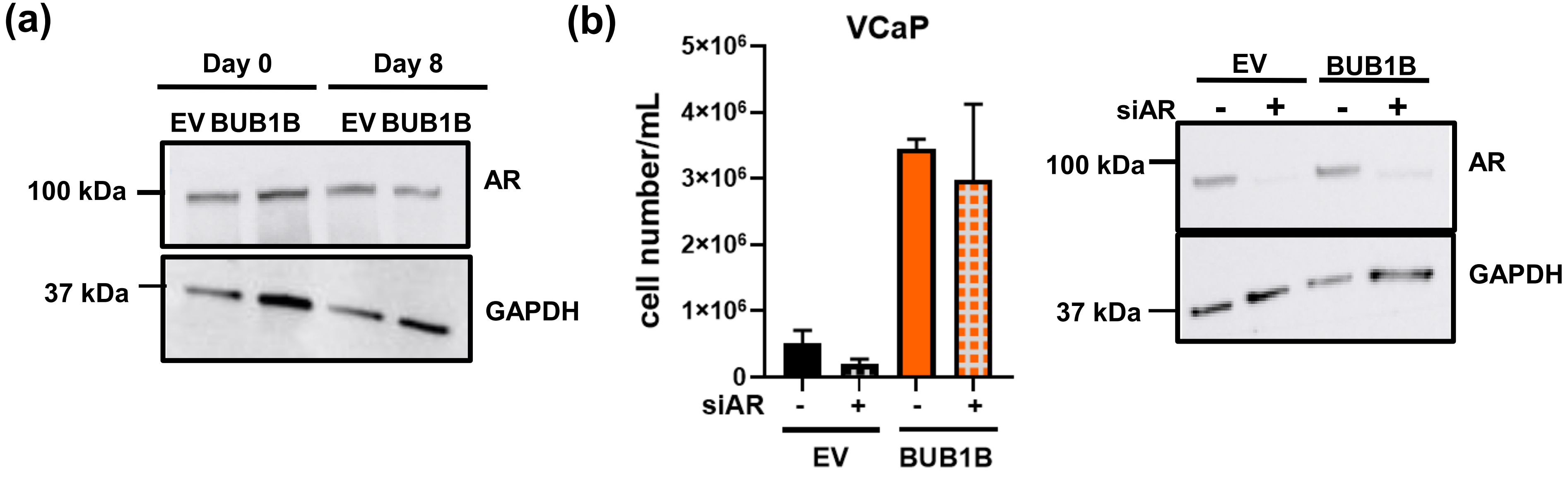


Supplementary Figure 4

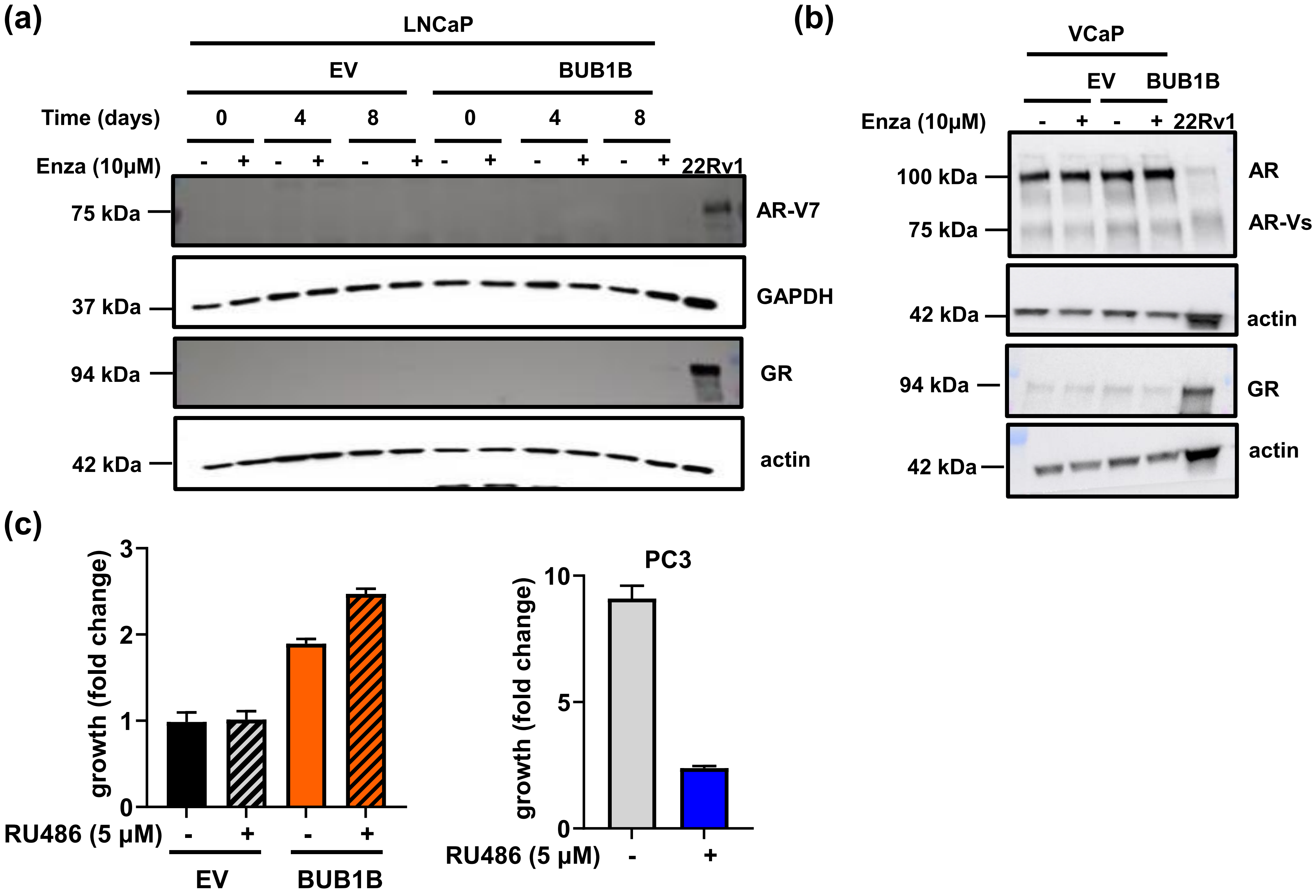


**Supplementary Figure 5**

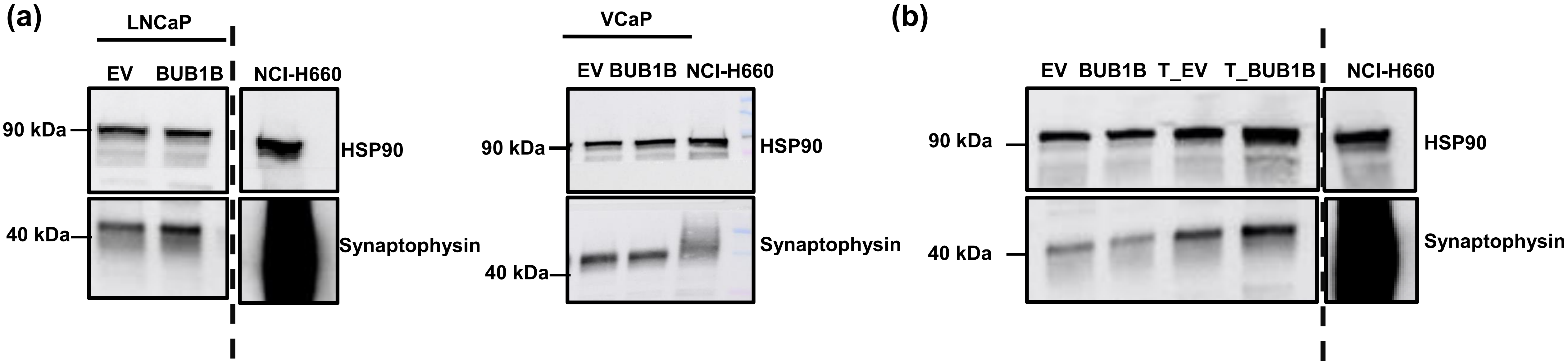


**Supplementary Figure 6**

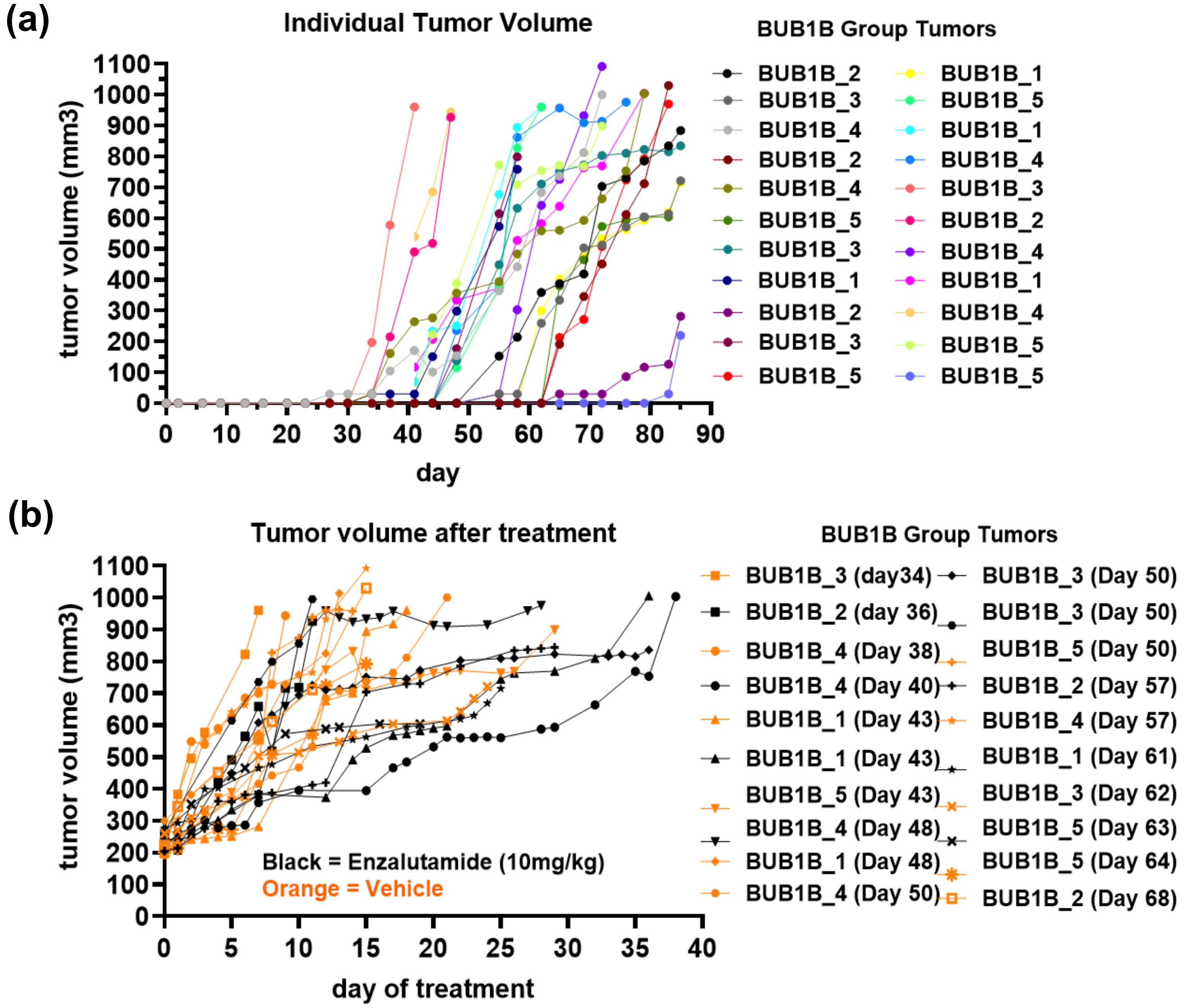


**Supplementary Figure 7**

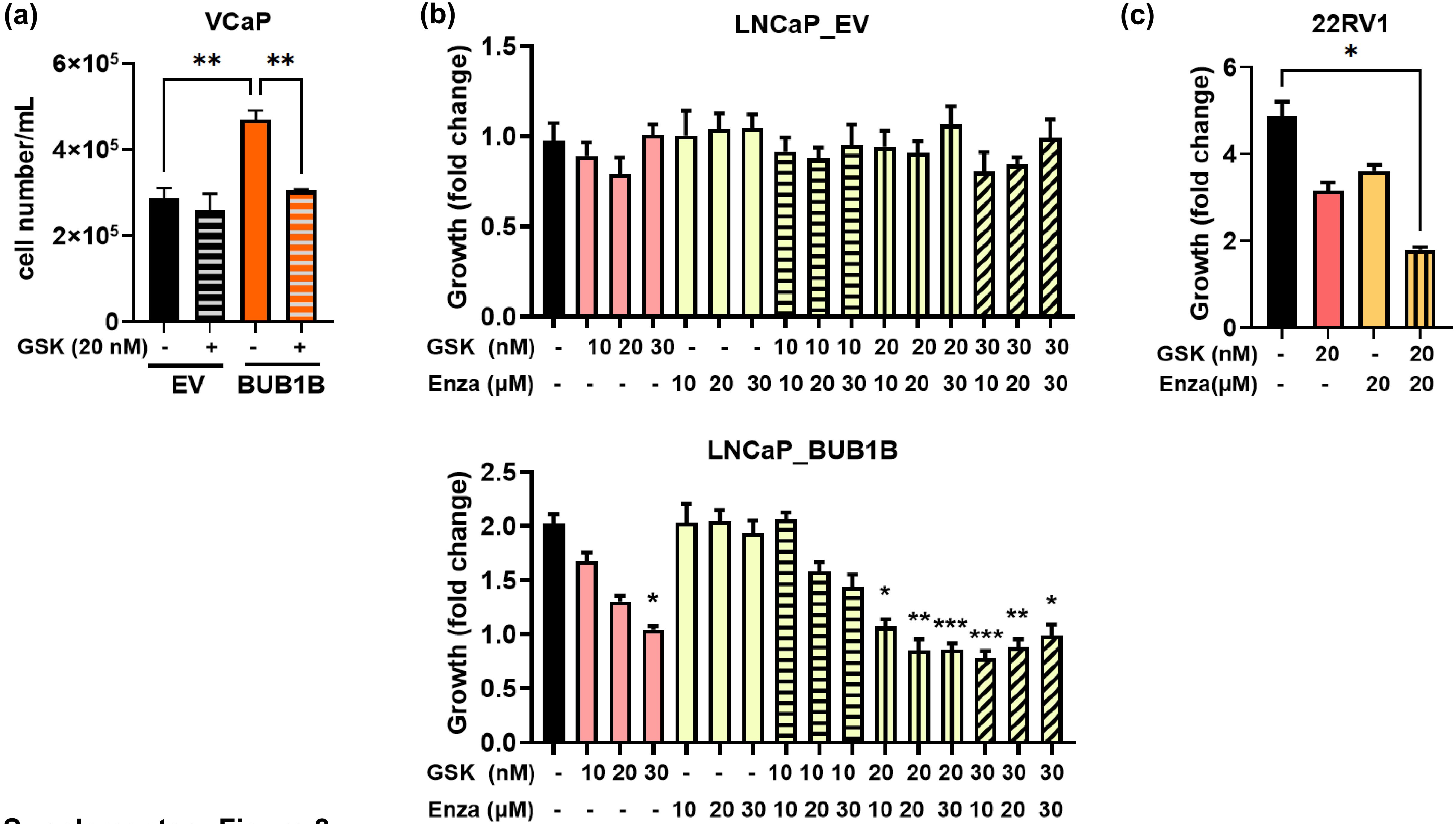


Supplementary Figure 8
